# Multiplexed and scalable cellular phenotyping toward the standardized three-dimensional human neuroanatomy

**DOI:** 10.1101/2022.11.23.517711

**Authors:** Tatsuya C. Murakami, Nathaniel Heintz

## Abstract

The advent of three-dimensional histological methods has advanced studies of cellular-resolution anatomy of the brain. The use of whole-mount staining and tissue clearing has advanced systems-level identification of cells underlying brain functions in mouse models. However, application of these methods to studies of human brains has been difficult due to their structural variability and the lack of standardized quantitative metrics. Here we report a rapid and scalable staining/imaging technique, termed mFISH3D, that enables single-cell-resolution imaging of mRNAs of more than ten genes in a large mammalian brain. To apply mFISH3D to postmortem human cerebral cortex, we have reconstructed morphogenic tracks of cortical growth, and used the tracks to provide a framework for quantitative assessment of cytoarchitecture. The workflow enabled the objective quantification of biological heterogeneity among cortical regions. We propose these techniques for standardization of 3D histology of the human cortex to provide reproducible measurements of cell-type-specific neuroanatomy.

## INTRODUCTION

The recent advances in tissue-clearing techniques and microscopy opened a door to three-dimensional (3D) single-cell-resolution neuroanatomy. Whole-mount tissue embedding and staining have allowed visualization of the intact cytoarchitecture of postmortem human brains (Hildebrand et al., 2019; Inoue et al., 2019; Ku et al., 2020; Lai et al., 2018; Morawski et al., 2018; Murray et al., 2015; Park et al., 2019; Pesce et al., 2022; Tainaka et al., 2018; Zhao et al., 2020). Concurrently, new optical configurations of microscopy, most notably light-sheet microscopy, were developed to image large transparent tissues at single-cell resolution (Chen et al., 2020; Dodt et al., 2007; Glaser et al., 2022, 2019; Huisken et al., 2004; Keller et al., 2008; Tomer et al., 2014; Voigt et al., 2019). The advent of these modern imaging techniques promised to advance our understanding of cellular mechanisms that support the functional diversities of the cerebral cortex. However, because of the limited capabilities of cell-type-specific labeling techniques in human brain samples, the application of these approaches remains restricted to regional surveys in thin tissue slices or non-cell-type-specific characterization of cytoarchitecture. Given these limitations, and the gyrencephalic and complex histological properties of the human cortex, there is presently no generally accepted scheme for computational deconvolution of 3D human imaging data that allows accurate, quantitative studies of the variation of human cell types across cortical regions, or in response to damage or disease.

Quantitative cell type characterization in the human cortex at high resolution is difficult due to the non-uniformity of cell densities and distribution that occur even within a single gyro-sulcal area. At the macroscopic level using magnetic resonance imaging (MRI), a variety of analytical workflows have been developed to deconvolve gyro-sulcal patterns of cortex (Jiang et al., 2021). One of the most successful techniques is cortical surface-based analysis and visualization with surface flattening (Fischl et al., 1999; Van Essen et al., 1998). Given the limited resolution of MRI, these cortical analytic workflows have not yet been applied for 3D cell-resolution studies. The advent of 3D single-cell-resolution imaging of the cleared human cortex has raised new challenges in the analysis of gyrencephalic brains. Surface-based analysis is helpful, but not sufficient unless we can associate each measurement in the volume with measurement of the surface. For example, the cortical thickness can be uniquely determined on any given point on the cortical surface (Fischl and Dale, 2000), but there is no explicit association between the local cell density of a certain point of gray matter with a specific point of the cortical surface. We can possibly make the association explicit by constructing connections that bridge the cortical surface and the boundary of gray-white matter as described previously (Amunts and Zilles, 2015; Kiwitz et al., 2020; Schleicher et al., 1999). The approach can be extended to 3D, but such abstract connections do not reflect the developmental history of the cortex, thus quantification based on such connections does not possess biological significance. To provide an analysis scheme that more closely reflects cortical development, we have developed a new approach in which connections are reconstructed based on the organization of cortical vasculature, which is thought to be a vestige of radial glial cell migration pathways used earlier in development (Paredes et al., 2018; Rakic, 2003). These connections, termed morphogenic tracks, follow the native flow of corticogenesis. The scheme enables quantitative evaluation of the impact of gyrification and enables segregation of the impact of gyrification for regional or pathological comparative analysis.

The ability to measure the number and distribution of specific cell types with this analytical framework can provide a step forward in studies of human brain anatomy. Historically, the comparative histology of human brain cortices has been examined with the aim of cortical parcellation through extensive observation of cytoarchitecture (Brodmann, 1909). Despite the utility of the technique for cortical parcellation, non-cell-type-specific cytoarchitectural observation provides little information about how the observed differences contribute to the functional diversity of the cortex, or how the disruption in cell composition relates to disease. To obtain mechanistic insight, a systematic evaluation of cell composition with molecular labeling is needed. To date, various 3D staining techniques have been proposed to identify the molecular properties of cells in human brain (Liebmann et al., 2016; Liu et al., 2016; Morawski et al., 2018; Murray et al., 2015; Nojima et al., 2017; Phillips et al., 2016; Tainaka et al., 2018; Tanaka et al., 2020; Zhao et al., 2020). However, a staining technique that satisfies scalability and multiplexability with enough simplicity has not been reported. Here, we report *m*ultiplexed *F*luorescence *I*n *S*itu *H*ybridization in *3D* (mFISH3D) for scalable molecular phenotyping of cells. Using a whole-mouse brain for demonstration, mFISH3D enabled visualization of mRNAs of ten genes, allowing the mapping of multiple cell types. We applied mFISH3D to quantify the composition of inhibitory neurons over the cerebral cortices of mouse brains, revealing a new relationship between satellite oligodendrocytes and somatostatin-expressing inhibitory neurons. Application of this approach to the human cerebral cortex, in combination with the use of morphogenic tracks for analysis, suggested that the variabilities of the cell density associated with gyrification may be explained by the flux of morphogenic tracks in gray matter. Furthermore, analysis of the prefrontal cortex and the secondary visual cortex demonstrated that morphogenic tracks can minimize confounding variables and enable quantitative comparison of cortical thickness and cell density. Using mFISH3D, we verified the higher density of excitatory neurons and oligodendrocytes in the secondary visual cortex than in the prefrontal cortex, and obtained evidence for a synchronous emergence of the excitatory neurons and oligodendrocytes. The proposed workflow is a new approach that enables quantitative comparative measurements of cell distribution for studies of regional or pathological variations in the human brains.

## RESULTS

### Fluorescent in situ hybridization with enhanced signal in a whole-mammalian brain

A variety of whole-mount staining technologies using fluorescent antibodies or in situ hybridization have been developed and employed for improved analysis of tissue anatomy (Belle et al., 2017; Cai et al., 2019; Hama et al., 2015; Murray et al., 2015; Renier et al., 2014; Susaki et al., 2020; Tanaka et al., 2017) and gene expression (Guo et al., 2019; Kumar et al., 2021; Sylwestrak et al., 2016; Tanaka et al., 2020). Despite these important advances, antibody penetration into tissue requires long incubation times and limits the scalability of the staining (Richardson et al., 2021). Volumetric staining using fluorescent in situ hybridization (FISH) takes advantage of the faster penetration of oligonucleotide probes and the location of target mRNAs in the cell soma to study gene expression in large tissues (Guo et al., 2019; Kumar et al., 2021; Sylwestrak et al., 2016; Tanaka et al., 2020). Notably, Kumar et al. demonstrated the staining of a mouse brain hemisphere in less than a week. However, in existing whole-mount in situ hybridization, it remains difficult to obtain a high signal-to-noise ratio across visible wavelengths using multiple FISH probes, and multi-round staining is limited by the fragility of mRNAs.

Published protocols for three-dimensional FISH share a common chemical strategy: delipidation or dehydration of the tissues, hybridization of primary probes against RNAs, amplification of fluorescent signal with hybridization chain reaction (HCR) (Choi et al., 2018; Dirks and Pierce, 2004), and tissue clearing (Guo et al., 2019, p.; Kumar et al., 2021; Sylwestrak et al., 2016; Tanaka et al., 2020). To optimize chemical compositions and reaction conditions in each step for the multi-color FISH of a whole-mammalian brain, we first formulated a 3D FISH protocol with minimal chemical components which published protocols have in common (**Figure S1A**), and termed it “the basic protocol”. We took a bottom-up approach to refine the basic protocol by modifying the chemical processes. We used an organic solvent in the tissue-clearing step to eliminate the impact of pH and the impact of salt concentration on hybridization. We excluded hydrogel-matrix-embedding to keep the optimization process simple. After confirmation that the basic protocol can produce a faint FISH signal, we further altered three steps of the basic protocol: tissue pretreatment, primary-probe hybridization, and HCR probe hybridization. In the tissue-pretreatment step, we included acid pretreatment and proteinase K digestion, which are known to be effective in thin tissue sections. Both pretreatments enhanced the signal-to-noise ratio in 2-mm-thick mouse brains (**Figures S1B and S1C**). In the primary-probe hybridization step, we explored a water-exclusion substance. While it is known that the water-exclusion substance such as dextran sulfate or polyethylene glycol assists hybridization efficiency (Knowles et al., 2011), we found that the large molecular-weight (MW) dextran sulfate (MW = 30,000) inhibited the probes from penetrating into the inside of the tissue. Replacing it with polyethylene glycol (MW of 8,000) achieved better probe penetration and improved hybridization efficiency (**Figure S1D**). We also used the same buffer for the HCR hybridization step as primary-probe hybridization. These alterations yielded a protocol that allowed the visualization of multiple gene transcripts across visible wavelengths in a whole-mount mouse brain (**Figure S1E** and **Supplementary Video 1**).

### Repeated cycles of FISH by heat-assisted hybridization cycling

The simplest approach to achieving multiplexed imaging is to repeat multiple rounds of staining/imaging and destaining cycles (**Figure S2A**). Though the approach has proven to be successful in a thin-tissue section such as in situ sequencing (Moses and Pachter, 2022), multi-round FISH staining in a whole-mammalian brain has not been achieved, presumably because the long incubation times can result in the degradation of mRNA. To overcome this problem, we took advantage of the increased stability of RNA:DNA hybrids (Kankia and Marky, 1999). We designed the oligonucleotide probes so that the primary probes would not result in off-target amplification of signals (**Supplementary Methods**). To prevent the carryover of the fluorescence from the previous cycles, we removed the primary probes and HCR probes by heat-assisted de-hybridization. Using this approach, we were able to detect the signal after six rounds of heat-assisted staining/destaining (**Figure S2B**). The carryover of the fluorescence from one cycle to the next was negligible (**Figure S2C**).

### Image processing and identification of molecular phenotypes of cells in a whole-mammalian brain

For single-cell-resolution imaging, we scanned the mouse brain using light-sheet microscopy with the pixel size of 1.3 μm × 1.3 μm at the optical sectioning step of 3.0 μm with 4 × 4 z-columns (**Supplementary Methods**). A single round of four-channel imaging of a whole mouse brain produces approximately 1 terabyte (TB) of data. Since we used a wide range of wavelengths from 488 nm to 785 nm as excitation light, chromatic aberrations could appear and cause focal shifts in the axial direction. One of the major sources of chromatic aberrations is illustrated in **Figure S2D**. After the detection of common feature points across channels, the shifts are corrected by linear registration (**Figures S2D and S2E**). Stitching of z-columns was also performed to compensate for the displacements of the motorized stages. We used Bigsitcher for both purposes (Hörl et al., 2019) (**Supplementary Methods**).

Another challenge in image processing is the registration of the images from different staining cycles. The registration has to take tissue deformation during staining cycles into account and thus must be non-linear. To obtain the registration with cellular-level accuracy, we included the signal of ribosomal RNA (*rRNA*) at each cycle. The registration of terabyte-scale datasets was previously reported (Wang et al., 2021), but the workflow was optimized for expanded tissue sections. While keeping the strategy of doing tile-wise non-linear registration after coarse global registration, we established the registration workflow with cellular precision for a large cleared tissue (**Figures S2F-S2I** and **Supplementary Methods**). To test the precision of the chromatic correction and the registration, we quantified the percentage of the cells which appear in the same position after the registration. Using a 3-mm-thick coronal mouse-brain block, we stained both *rRNA* and mRNA of neuropeptide Y (*Npy*). After the two rounds of staining and imaging cycles, *Npy*-expressing cells were segmented and subjected to analysis. We found that 97.4% of the segmented cells appeared in the same position with the intersection over union (IoU) threshold of 0.4 (**Figure S2I**).

By targeting distinct sets of mRNAs in each cycle, we can reveal the molecular identities of cells in a whole brain. We term this method mFISH3D. We targeted in total 10 gene transcripts, *rRNA*, vasoactive intestinal peptide (*Vip*), solute carrier family 17 member 7 (*Slc17a7*), proteolipid protein 1 (*Plp1*), cannabinoid receptor 1 (*Cnr1*), *Npy*, glutamate decarboxylase 1 (*Gad1*), somatostatin (*Sst*), cholecystokinin (*Cck*), and tyrosine hydroxylase (*Th*) (**Figure 1**) using mFISH3D and co-stained with alpha-smooth-muscle actin (α-SMA) using immunostaining (**Supplementary Methods**).

**Figure 1.**
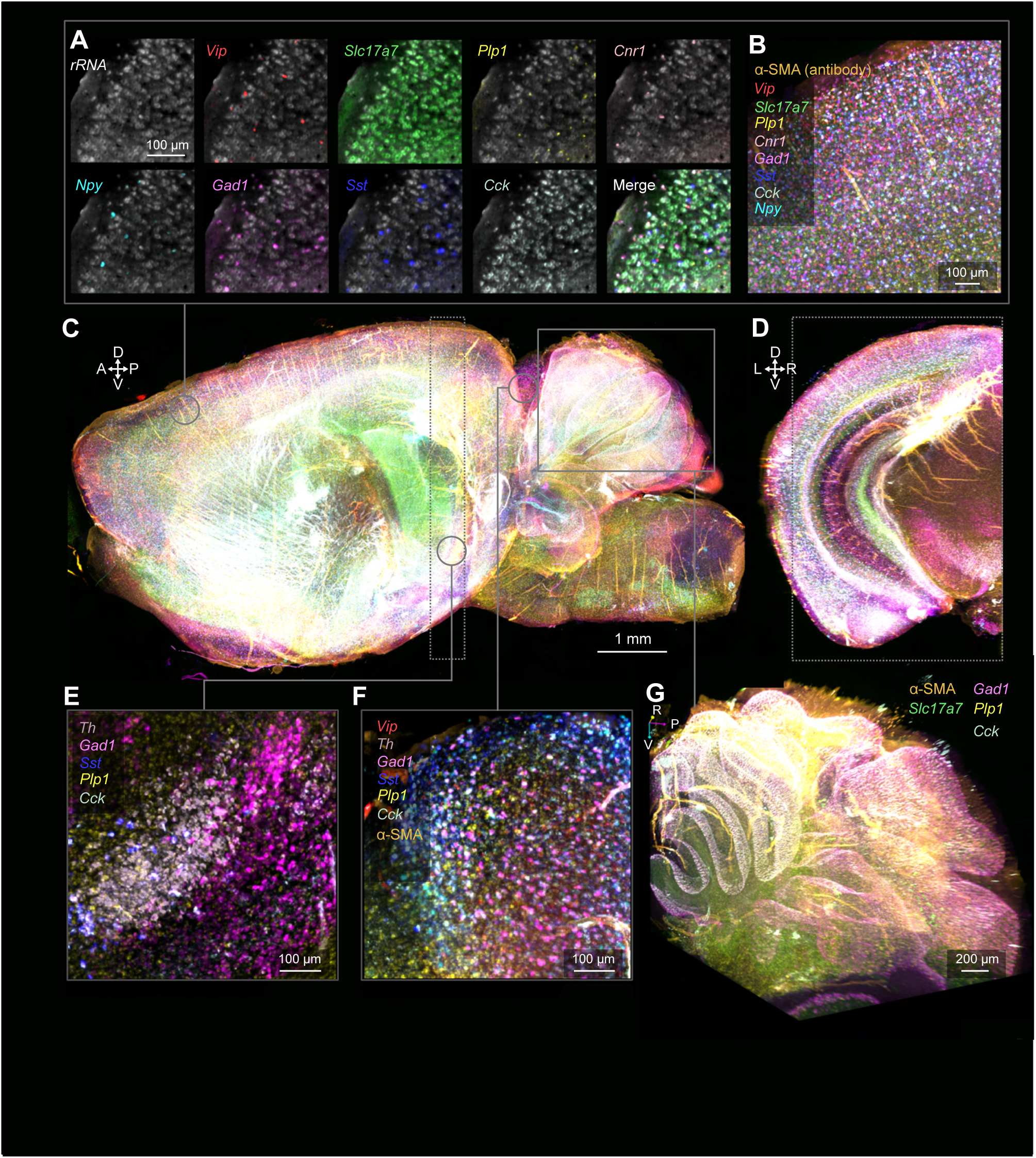
Whole-brain multiplexed fluorescent in situ hybridization (mFISH3D) (**A**) The magnified view of the mouse-hemisphere brain stained with mFISH3D. Part of the cerebral cortex is shown. Each panel includes signals of *rRNA* and mRNA of a gene. Six rounds of imaging were registered in one space. mFISH3D was performed against *rRNA*, *Vip*, *Slc17a7*, *Plp1*, *Cnr1*, *Npy*, *Gad1*, *Sst*, *Cck*, *Th*, and co-stained with anti-alpha-smooth muscle actin antibody (αSMA). (**B**) The maximum-intensity projection of the cortex. The volume was projected over 150 μm along the left-right (LR) axis. (**C**) The sagittal view of the volume rendering of the 10-gene-expression pattern in the hemisphere. The gene-expression patterns except *rRNA* are shown. The area marked by the dotted rectangle was used for **D**. (**D**) The coronal view of the volume rendering of the hemisphere, marked by the dotted rectangle in **C**. (**E**) The maximum-intensity projection of the region around the substantia nigra. The volume was projected over 150 μm along the LR axis. (**F**) The maximum-intensity projection of the region around the colliculus. The volume was projected over 150 μm along the LR axis. (**G**) The perspective volume rendering of the cerebellum, marked by the solid rectangle in **C** was used. A: anterior, P: posterior, D: dorsal, V: ventral, L: left, R: right.

### Cortex-wide mapping of subtypes of inhibitory neurons

We performed mFISH3D on three whole brains of adult wild-type C57BL/6J mice by targeting *rRNA* and mRNAs of *Gad1*, parvalbumin (*Pvalb*), *Sst*, *Vip*, *Npy,* and *Plp1* (**Supplementary Video 2**). The stained cell bodies were segmented over the entire cerebral cortices (**Figure 2A** and **Supplementary Methods**). The cells were annotated as inhibitory neurons based on the expression of *Gad1*. The inhibitory neurons were further classified into *Vip*, *Sst,* or *Pvalb*-expressing neurons (hereafter *Vip*+, *Sst*+, and *Pvalb*+) based on the expression of the three canonical marker genes, which are differentially expressed in functionally distinct interneurons (Tremblay et al., 2016). The inhibitory neurons without expression of *Vip*, *Sst,* and *Pvalb* were labeled as “Other”. The expression of *Npy* can also mark subtypes of inhibitory neurons, but it is known that subsets of *Vip*+, *Sst*+, and *Pvalb*+ cells each express *Npy* (*Npy*+) (Karagiannis et al., 2009; Tremblay et al., 2016). Using this fact, we further subclassified the four major classes of inhibitory neurons (*Vip*+, *Sst*+, *Pvalb*+, and Other) based on the expression of *Npy*. To confirm whether the distribution of cells was consistent with previous reports, we compared our counts of cells to the counts from the study by Kim et al. (Kim et al., 2017) and obtained a high correlation (**Figure S3A**). After registering the brains to the Allen Brain Atlas Common Coordinate Framework v3 (CCFv3) (Wang et al., 2020), we revealed the number of each subtype of inhibitory neuron (**Figure S3B**). We analyzed the ratio of *Npy*+ cells within the four major subtypes (**Figures 2B** and **2C**) and found that the ratios of *Npy*+ largely depended on the cell types, regions, and layers in the cortices.

**Figure 2.**
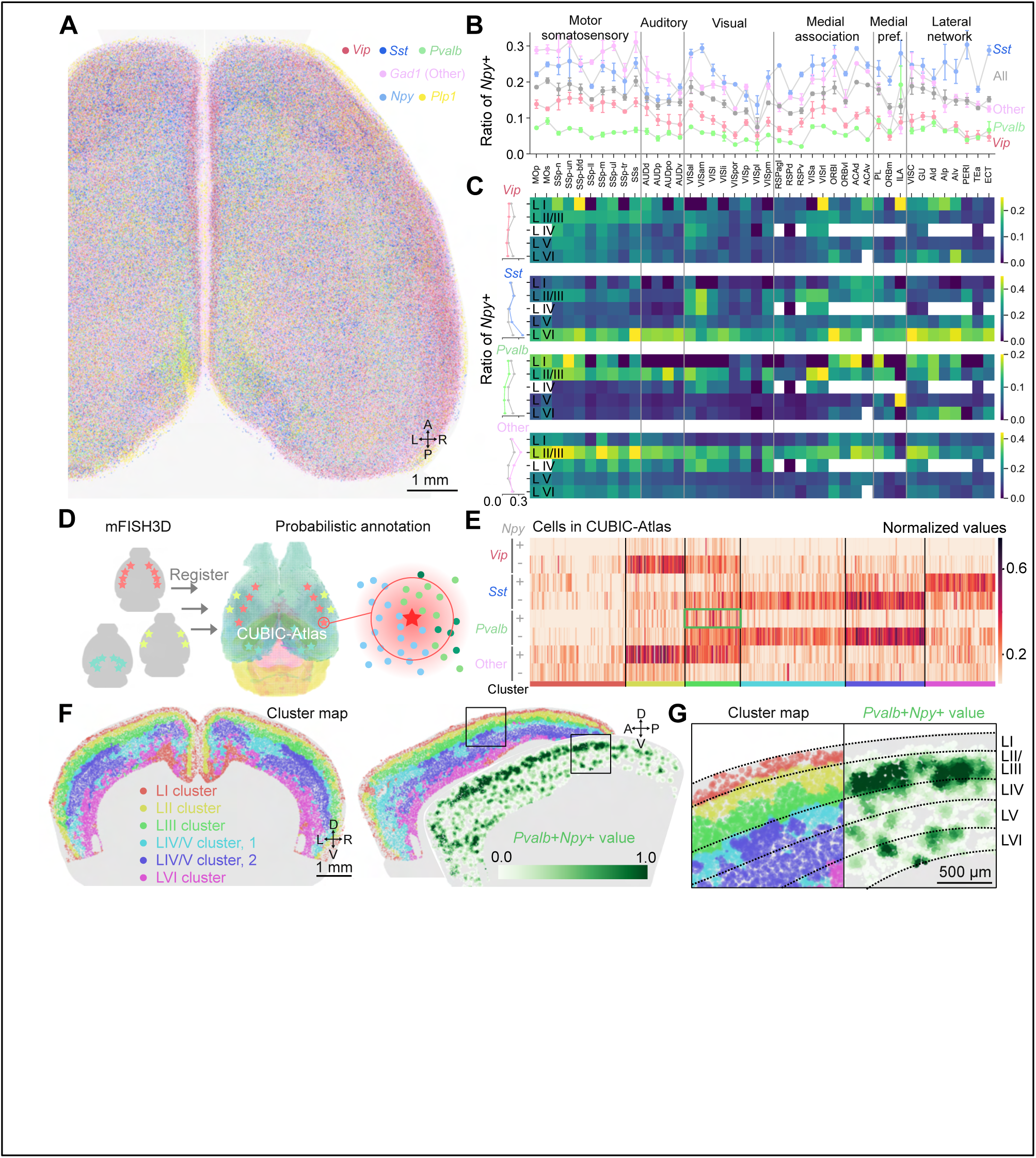
Cortex-wide mapping and classification of inhibitory neurons using CUBIC-Atlas led the discovery of the *Pvalb*+*Npy*+ subpopulation. (**A**) The marker mRNAs of inhibitory neurons (*Gad1*, *Vip*, *Sst*, *Pvalb*, *Npy*) were stained with mFISH3D together with the marker mRNA of oligodendrocytes (*Plp1*). The cell bodies were segmented and the centroids were visualized. After the registration to Allen Brain Atlas Common Coordinate Framework (CCFv3), the cortical cells of one representative brain were visualized. (**B**) The *Gad1*+ cells were classified into four groups (*Vip*+, *Sst*+, *Pvalb*+, and Other). The four groups were further classified based on the expression of *Npy*. The percentages of *Npy*+ in the four classes were calculated and their regional variations are shown over isocortices (N=3). The annotations of the regions were obtained from CCFv3. The functional classification of the regions was obtained from a previous study (Kim et al. 2017). The values are mean ± standard deviation (s.d.). Medial pref.: Medial prefrontal. (**C**) The percentages of *Npy*+ in the four classes are shown by cortical layers in the heatmaps. The annotations of the layers were obtained from CCFv3. The white blanks indicate that the regions did not have the corresponding layer(s) in CCFv3. The averaged percentages by layer are shown on the left. The gray line indicates all inhibitory neurons. (**D**) The scheme of single-cell-resolution analysis using CUBIC-Atlas. (**E**) The heatmap visualization of the normalized values after the probabilistic annotation of the CUBIC-Atlas. After k-means clustering, the annotated cells were aligned according to the allocated clusters. (**F**) The spatial mapping of the clusters in a coronal slice view (left) and sagittal slice view (right). In the sagittal view, the normalized values of *Pvalb*+*Npy*+ are also visualized. (**G**) The magnified sagittal views of cluster map and *Pvalb*+*Npy*+ values. The annotations of the layers are from CCFv3. The rectangular areas from **F** are shown. A: anterior, P: posterior, D: dorsal, V: ventral, L: left, R: right.

While the region-wise analysis can describe the cellular composition over functionally distinct regions, the downside is that the analysis ignores the importance of sub-localization within the annotated regions. For analysis at higher resolution, we used CUBIC-Atlas, a single-cell resolution atlas (Mano et al., 2021; Murakami et al., 2018). After the registration and probabilistic annotation of the CUBIC-Atlas, we performed a clustering analysis to uncover cellular populations with unique distribution patterns (**Figures 2D** and **2E**). We found that 6 clusters were localized according to the laminar pattern of the cortex (**Figure 2F**). Two clusters, shown in cyan and blue, localized to layers IV and V. The cyan cluster had a lower content of *Sst*+ and *Pvalb*+ cells than the blue cluster, consistent with previous findings (Kim et al., 2017). We also found that the green cluster (LIII cluster) exclusively localized to layer III (**Figure 2G**). This cluster was unique in terms of its high content of *Pvalb*+*Npy*+ cells. Although the *Pvalb*+*Npy*+ cells were a minor neuronal population (**Figure S3B**), their predominant localization within layer III may indicate a functionally distinct role from other *Pvalb*+ cells.

### Satellite Oligodendrocytes preferably colocalize with *Sst*+ cells

Satellite oligodendrocytes are present throughout the cerebral cortex. In contrast to oligodendrocytes that form the myelin sheath from distal positions, their cell bodies are found in close apposition to neuronal soma. Although the function of the satellite oligodendrocytes is not fully clear, they are thought to support the metabolism of the adjacent cells (Takasaki et al., 2010). Given the diversity of neuronal types in the cerebral cortex, we asked whether satellite oligodendrocytes maintain preferential associations with specific classes of cortical interneurons. *Plp1* is one of the canonical marker genes for oligodendrocytes. After visually confirming that a portion of the oligodendrocytes appears next to inhibitory neurons (**Figure 3A**), we segmented *Plp1*+ cells and extracted *Plp1*+ cells which overlapped with the segmentation of inhibitory neurons. These candidate satellite oligodendrocytes were then further analyzed by measuring two probabilities. Theoretical probabilities were calculated based on the local density of the subtypes of inhibitory neurons around the candidate oligodendrocytes. Empirical probabilities were calculated from the frequency of the subtypes of inhibitory neurons being adjacent to the candidate oligodendrocytes. The odds ratio was calculated for each candidate oligodendrocyte (**Figure 3B** and **Supplementary Methods**). The cortical-wide quantification revealed that the satellite oligodendrocytes were more likely to be bundled together with *Sst*+ cells than other types of inhibitory neurons (**Figures 3C** **and S3C**) and that this association was more frequently observed in deep layers of the cortex (**Figure 3D** and **S3D**). While the biased myelination of an interneuron subtype has been observed previously (Zonouzi et al., 2019), the preference of satellite oligodendrocytes for the *Sst*+ interneurons has not been reported. The preferential association of *Sst*+ cells and satellite oligodendrocytes may be due to their influence on oligodendrogenesis cells during brain development (Voronova et al., 2017), although further research is needed to understand the mechanisms responsible for this association.

**Figure 3.**
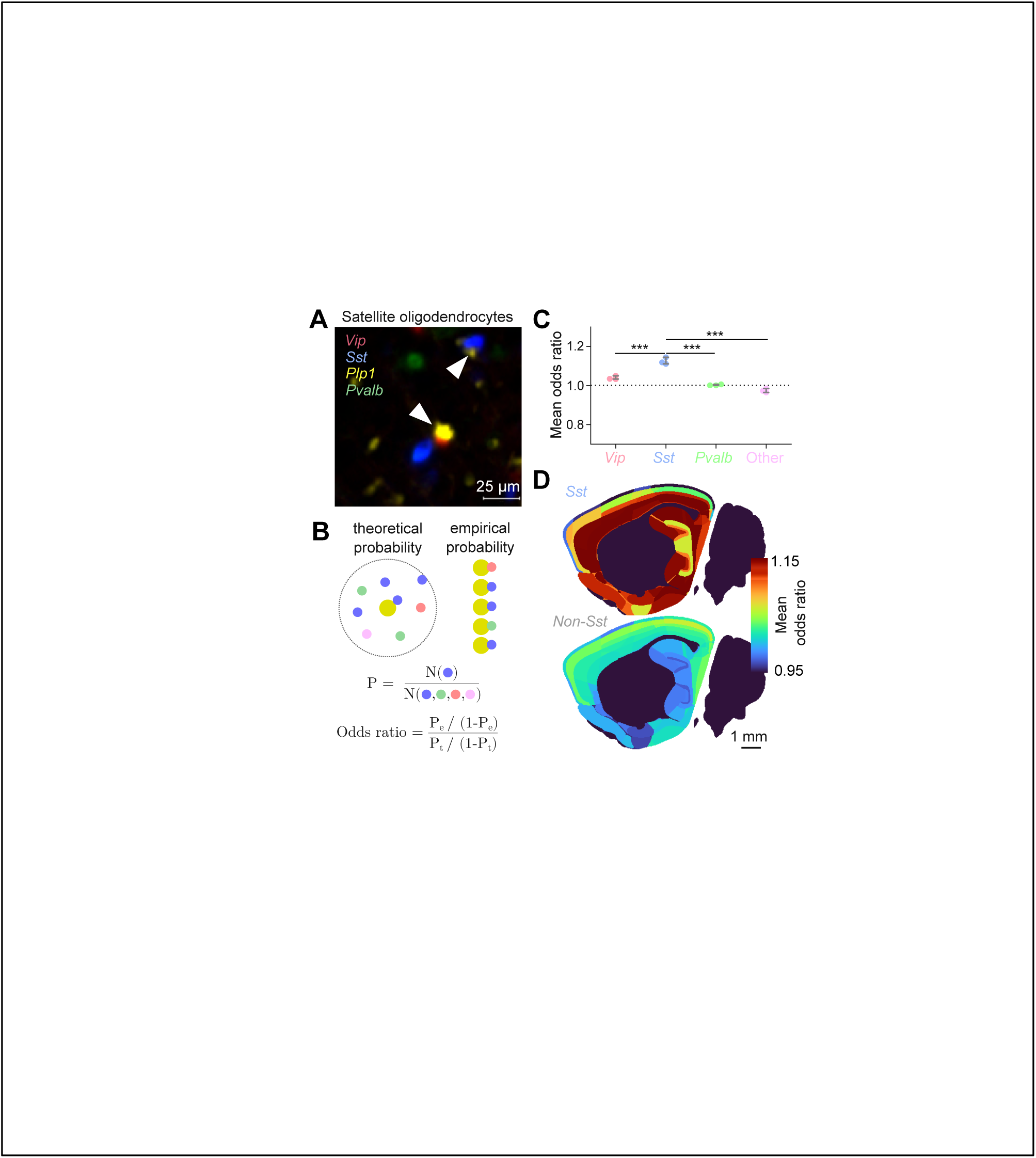
Interaction analysis between inhibitory neurons and oligodendrocytes revealed biased colocalization of *Sst*+ cells and satellite oligodendrocytes. (**A**) Representative image of the candidates of satellite oligodendrocytes. The possible satellite oligodendrocytes are indicated with arrowheads. (**B**) The derivations of theoretical probability (*P_t_*), empirical probabilities (*P_e_*), and odds ratio. The *P_t_* was calculated from surrounding cell densities while *P_e_* was calculated from the number of adjacent cells to the oligodendrocytes (**Supplementary Methods**). (**C**) Mean odds ratios by subtypes of inhibitory neurons in the cerebral cortices. ***P value < 0.001, one-way ANOVA with Tukey’s post hoc test for multiple comparisons (N=3). The error bars are s.d. (**D**) Sagittal maps of the mean odds ratio. The mean odds ratios for *Sst*+ cells are shown at the top. *Vip+, Pvalb+,* and Other cells were grouped together and their mean odds ratios are shown at the bottom as *Non-Sst*. The mean odds ratios in the cerebral cortex, olfactory areas, and hippocampal formation are visualized.

### Challenges in the quantification of cell compositions in gyrencephalic brains

While mFISH3D is a powerful tool for the spatial profiling of cells in a mammalian brain, the folded anatomy of the cerebral cortex of primate brains often hinders point-to-point quantitative comparison of regional cellular composition. This challenge of the gyrencephalic brain is further compromised by the fact that measurements such as cell number, cortical thickness, and cell density can be easily affected by small deviations in observation. For example, depending on the position in the gyrus and the plane chosen for analysis, measurements of cortical thickness can vary substantially in each laminar (**Figure 4A**) (Amunts et al., 2013). Furthermore, given the size of the human brain and local variations across individuals, it is often difficult to sample precisely a specific sulcus in a given brain region. Finally, even within a given gyrus/sulcus, cell density can vary depending on location with respect to the crown or fundus. To understand the impact of these variables on cell density, we simulated how changing angles in tissue sectioning affects the cell numbers in layers I-IV. Using SytoX^TM^ Green nuclear staining, we performed a three-dimensional scan of the ∼4×4×3 mm^3^ human cerebral cortex. Several 50-μm-thick tissue slices were digitally simulated from the volumetric image. We found up to ∼50% change in cell number within the range of 50° and we observed ∼20% change in cell number within the range of 20° (**Figures 4B**, **S4A**, and **S4B**). This variability is a major confounding factor for the analysis of human neuroanatomy at cell resolution.

**Figure 4.**
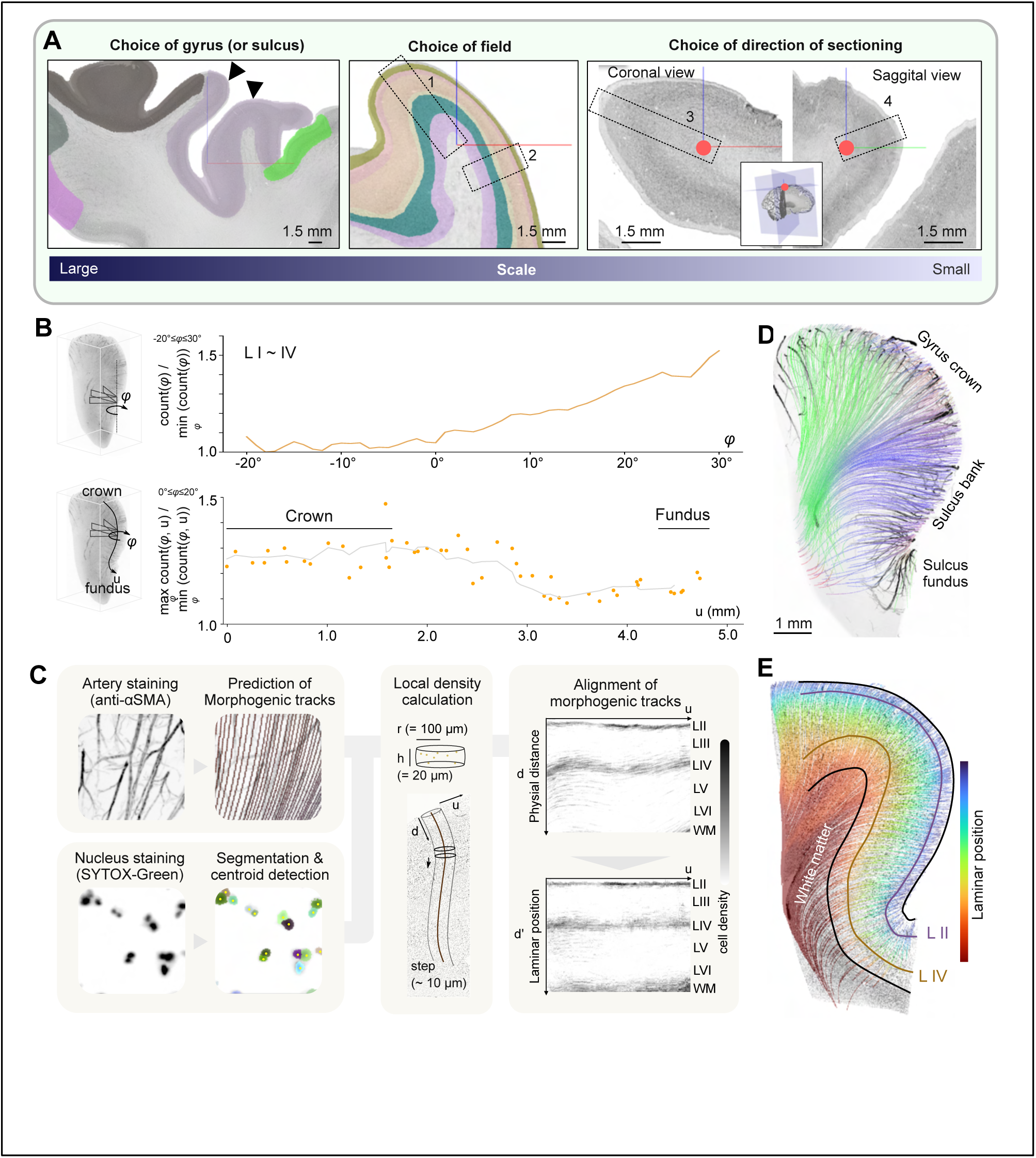
Reconstruction of morphogenic tracks in a human cortex. (**A**) Challenges in human neurohistology. Left: heterogeneity among gyri/sulci in a region. The arrowheads indicate different gyri. Distinct cortical regions are marked by colors. Middle: heterogeneity within a gyrus/sulcus. The fields labeled 1 and 2 show distinct laminar thicknesses. The layers are marked by colors. Right: impact of the choice of direction of sectioning. The red dots indicate identical positions. The volumetric image is indicated in the inset. The fields labeled 3 and 4 show distinct laminar thicknesses. The pictures and regional or layer annotations were obtained from the BigBrain project (Amunts et al. 2013) and the pictures were modified for the purpose of illustration. (**B**) Simulation of the variabilities that can be caused by conventional tissue sectioning. The virtual tissue slices with 500-μm width and 50-μm thickness were simulated from the imaging volume and their variabilities in cell number in layers I∼IV were quantified. The top indicates the impact of the changes in the angle of sectioning on the cell number. The sectioning angles were tested in the 50° range. The values normalized by the minimum counts are shown. The bottom shows how the variabilities change depending on the position relative to the gyrus. The sectioning angles were tested in the 20° range. We measured the ratio of the maximum count in the 20° range and the minimum count in the 20° range at each position. The moving averages are indicated by the gray line. (**C**) The workflow of predicting morphogenic tracks and calculation of the local cell densities. The alignments of morphogenic tracks were performed so that cell populations in the same laminar position were comparable (**Supplementary Methods**). (**D**) The reconstructed morphogenic tracks in the human prefrontal cortex. The tracks were color-coded based on the orientation of the local vectors. The signal of immunostaining of αSMA is overlaid and shown in grayscale. The 1-mm-thick volume is reconstructed for the visualization purpose. (**E**) The visualization of the laminar positions. The tracks were color-coded based on the laminar positions. The manual annotations of the cortical surface, layer II, layer IV, and boundary of gray and white matter are shown as landmark references.

### Reconstruction of morphogenic tracks in a human cerebral cortex

To eliminate this artificial variability, we propose an alternative approach that we believe reflects morphogenic flows of cells during cortical development. During corticogenesis, cellular migration, differentiation, angiogenesis, and axogenesis occur according to programmed biological guidance with a roughly consistent flow. For example, the radial glial cells form radial scaffolds for axial cellular migration (Rakic, 2003), arteries form a radial network that traverses the cortex (Kirst et al., 2020; Miyawaki et al., 2020; Todorov et al., 2020), and axons of neurons with long-range projections are guided in a radial direction then converge in white matter (Cajal, 1911). The morphogenic flows from those events are presumably interrelated. Notably, arteries have been reported to interact with radial glial cells whilst assisting cell migration and differentiation (Paredes et al., 2018). Given the impact of the arteries during morphogenesis, we decided to use the paths of arteries to predict radial morphogenic flows of the human cortex. The paths of arteries stained with anti-alpha-smooth muscle actin (αSMA) antibody, were used to generate a vector field where each vector indicates the local directions of morphogenic flows (**Figure S4C**). We then derived morphogenic tracks from the vector fields (**Figures 4C**, **4D**, and **Supplementary Video 3**). The morphogenic tracks were reconstructed from the surface of the cortex toward the white matter following the vector fields. As the starting points of the tracks, termed seeds, we chose the positions on the surface where the outermost cells reside (**Supplementary Methods**). We set the positions of the seeds to be the 0-μm position of the morphogenic tracks. Since cortical thickness varies depending on the positions relative to a gyrus, physical distance from the seeds is not an accurate predictor of the position of cortical layers. We thus developed the analysis workflow to align the positions of the cortical layers of the morphogenic tracks (**Figure 4C** and **Supplementary Methods**). In the workflow, we first detect the centroids of the nuclei in the cortex (**Supplementary Video 4**). We profiled the transition of the local density of nuclei along a morphogenic track and did the same analysis over all morphogenic tracks. Picking one representative morphogenic track as a reference, we aligned each morphogenic track to the reference morphogenic track. The alignment allows us to determine the positional correspondence among the morphogenic tracks. The accuracy of the alignment was examined by confirming that the aligned positions matched the ground-truth cortical layers (**Figures 4E** **and S4D**).

### Analysis of transitions of laminar composition over a gyrus

For explanatory purposes and data visualization, we introduced a coordinate system and terms as illustrated in **Figure S5A**. As is widely done in the visualization of MRI data (Fischl et al., 1999; Van Essen et al., 1998), we flattened the surface of the cortices and introduce this transformed coordinate as surface-based coordinates. Since the seeds of the morphogenic tracks can be shown in surface-based coordinates and one seed uniquely generates one morphogenic track, we used surface-based coordinates to display values related to morphogenic tracks. Using three tissue pieces from the prefrontal cortex of one donor, we examined the thickness and cell number in the laminar for each morphogenic track and visualized them in surface-based coordinates (**Figure 5A**). The profiling indicated that the cell numbers and thicknesses of layers vary depending on whether the morphogenic tracks begin from the gyrus or the sulcus. In both cell numbers and thicknesses of layers, the values were higher in the upper layer in a sulcal fundus than in a gyral crown, while the values were lower in the deep layers in a sulcal fundus than in a gyral crown. This observation is consistent with the previous findings that the thickness of upper layers I and II are known to be larger in the sulcal fundus and smaller in the gyral crown, while deep layers V and VI are thinner in the sulcal fundus and thicker in the gyral crown (Brodmann 1909; Wagstyl et al. 2020). The high correlation between the thickness and the number of cells (Pearson’s r = 0.96) implies we can roughly predict the number of cells along a morphogenic track given the thickness of the layers (**Figure 5B**). This fact motivated us to investigate how the thickness of the layer may be determined. We examined whether the thickness of the deeper cortical layers is enough to predict the thickness of the upper cortical layers. We found there is no clear correlation between the thickness of the deeper cortical layers and the more superficial cortical layers (Pearson’s r = -0.06) (**Figure 5C**). This may be due to non-uniform cortical growth during gyrification (Ronan and Fletcher, 2015).

**Figure 5.**
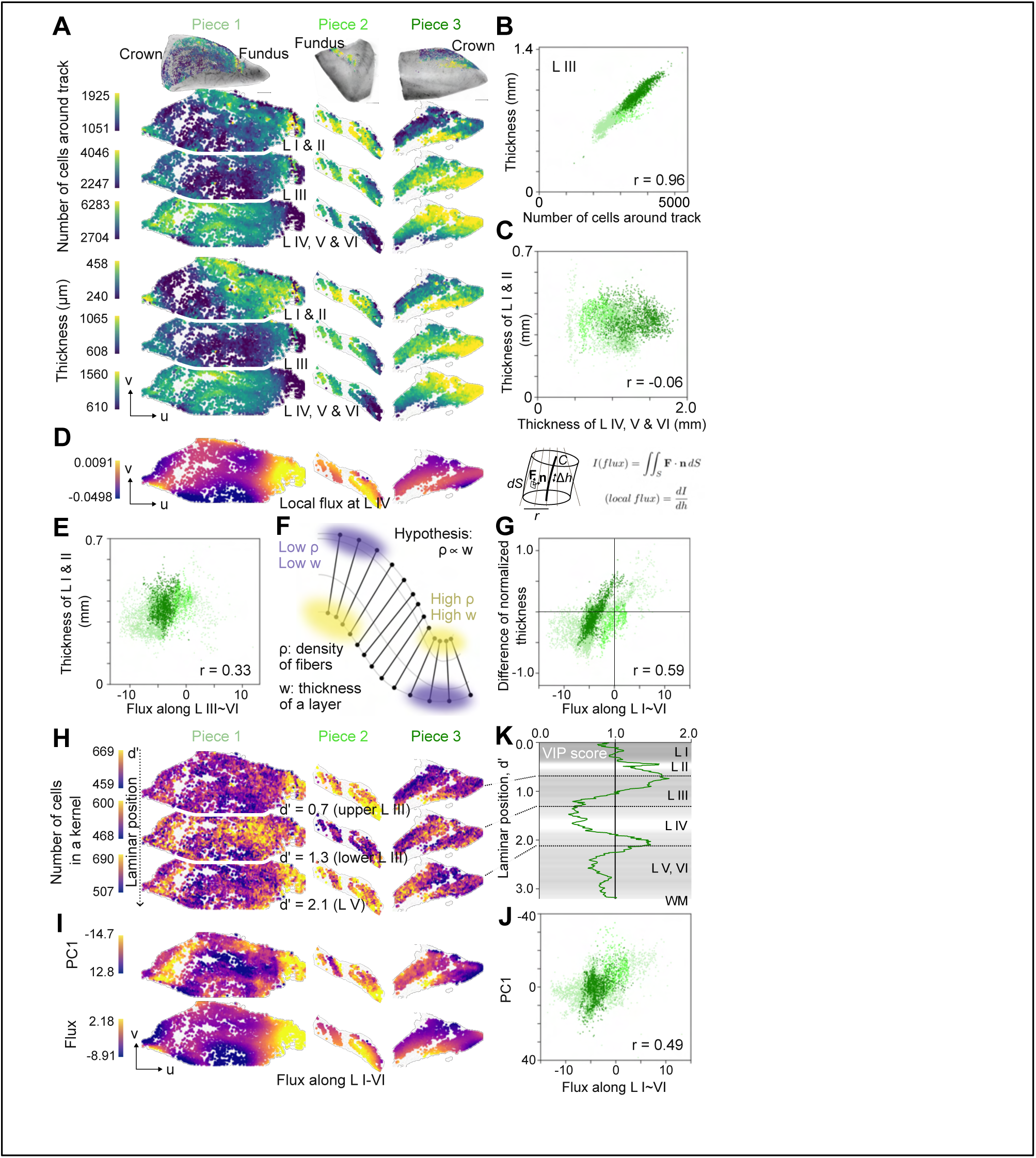
Analysis of the thickness of layers and cell density in the prefrontal cortex and their correlation with flux. (**A**) The number of cells and the thickness for each track by layers. Three pieces of the prefrontal cortex from one donor are visualized in surface-based coordinates. The values were color-coded. The volume renderings of the pieces are shown at the top. (**B**) Scatterplot of the number of cells around a track and the thickness at layer III. (**C**) Scatter plot of the thickness of deep layers (sum of layers IV, V, and VI) and the thickness of upper layers (sum of layers I and II). (**D**) Left: the local flux at layer IV in surface-based coordinates. Right: the definition of flux and local flux in this study. (**E**) Scatter plot of the flux along layers III∼VI and the thickness of layers I and II. (**F**) Illustration of the model. The model hypothesizes the higher fiber density (ρ) ends up in the higher thickness (w) of the layers. The model assumes the density of the fibers is proportional to cortical thickness. (**G**) Scatter plot of the flux along layers I∼VI and the difference of normalized thickness. The thickness of the upper layer (layers I and II) and the deep layers (layers IV∼VI) are measured and the values were separately normalized by the median values. The differences between the two normalized values were calculated for each morphogenic track. (**H**) The visualization of local cell density at each laminar position. The values were color-coded according to the bars at the left. (**I**) The visualization of the principal component 1 (PC1) and the flux along layers I-VI. (**J**) Correlation plot of the PC1 and the flux along layers I-IV. (**K**) The variable importance of projection (VIP) score of partial least square regression. The laminar positions with higher VIP scores can be assumed to have a higher correlation with the flux. The averaged cell density with a gray color code is shown in the background. Note that both flux (1/µm^2^) and local flux (1/µm) were scaled by an arbitrary constant in this figure. L: layer, WM: white matter.

To take the impact of gyrification into account, we introduce the concept of flux and local flux (**Figure 5D**, right). Flux can be measured for each morphogenic track as the relative amount of fibers (i.e. arteries) that the surrounding area of the morphogenic track gains or loses. Because the flux can be impacted by the thickness of the cortex, we also introduce the term local flux, which reflects the flux per unit thickness to exclude the impact of the thickness from the analysis. In general, the positive flux or local flux means the flow is converging, as seen in sulci, while the negative flux or local flux means the flow is diverging, as seen in gyri. The local fluxes at layer IV are visualized on the left of **Figure 5D**. The sulci have higher values than gyri in general. Interestingly, the flux of the deeper layers shows a moderate correlation (Pearson’s r = 0.33) with the thickness of the upper cortical layers (**Figure 5E**). This finding led us to formulate a model that can potentially explain the source of the different cortical thicknesses over a gyrus. The model assumes that the thickness of the layer is proportional to the density of fibers (**Figure 5F**). Note we used the term “fibers” in a different sense from the “tracks”. The fibers have biological entities such as arteries or radial glial fibers. We proposed the model based on the assumption that the higher density of fibers leads to a higher supply of nutrients and metabolites, thus leading to enhanced growth and cortical thickness. It is important to note that this model does not depend on the distribution pattern of cells in layers.

Because the local density of the arteries is difficult to estimate due to their sparsity, we used the fluxes along layers I∼VI, which corresponds to the gain of the arteries in layers I∼VI, and compared them with the difference in normalized thicknesses between the upper layers and deep layers (**Figure 5G**). According to the model, the higher the flux, the higher the difference, and the relation of these values would be linear. Consistent with the model, we see a high linear correlation between these values (Pearson’s r = 0.59). It will be interesting to test the generality of the model by expanding the imaging volume. This hypothesis will bring a microscopic perspective to the cortical gyrification theory (Ronan and Fletcher, 2015; Van Essen, 1997).

### Correlation between flux and cell densities

In addition to the thickness of the cortex, the densities of cells within the cortical laminar are also important and thought to impact brain function. For example, changes in cortical thickness and cell density have been reported in several neurological disorders (Cotter et al., 2001; Hanford et al., 2016; Thu et al., 2010; Zhao et al., 2022). However, little is known about the variability of cell densities within a gyrus/sulcus and how it is determined. It is also important to analyze at a finer resolution than the laminar structures because some of the cell types do not follow the canonical laminar labeling. For this purpose, we used aligned distance (*d’*) as a metric to indicate cell positions as a finer alternative to layers. We hereafter call this aligned distance a “laminar position”. We set the surface of the cortices to be the beginning of the laminar position (*d*’=0) (**Figure S5B**). Visualization of the cell densities in surface-based coordinates indicated that the density pattern was not constant over gyri/sulci (**Figure 5H**). Importantly, the pattern of the cell density is dependent on the laminar position. We investigated the major factor to affect the pattern of cell density. We performed a principal component analysis (PCA) to obtain a summarized value on a morphogenic track and observed the spatial pattern of the value. Interestingly, the first principal component showed a spatial pattern that correlates with the flux along the layers (Pearson’s r = 0.49) (**Figures 5I** and **5J**). This indicates that the flux is one of the major sources of the variability in cell density. This fact has practical importance because we can control the variability of cell density within a gyrus/sulcus by using flux. This utility of the flux was fully harnessed in the comparative analysis of the cortical regions which we demonstrate below. We also analyzed which laminar positions showed a strong correlation to the flux. We applied a Partial Least Square (PLS) regression model so that the model can predict flux from cell densities, and measured the Variable Importance in Projection (VIP) score to know which laminar positions display a high correlation to the flux (**Figure 5K**). We found that the correlation with flux is stronger in the upper layer III and layer V than in the deep layer III and layer VI. It is important to note that the flux is not the only source of the variability in cell density, especially in lower layer III and layer VI. Another important finding is that both upper layer III, which is close to the upper layer, and layer V, which is the deeper layer, have positive correlations with flux (**Figures 5H**, **5I**, and **5K**). This clearly differs from the cortical thickness, where the upper layer positively correlated with flux and the deep layer negatively correlated with flux (**Figures 5A** and **5I**). These observations imply that the cell density is not regulated in the same way as thickness. The mechanism of this irregular correlation between the flux and cell densities will require further investigation. With the assistance of morphogenic tracks, we can quantify the biological heterogeneity while minimizing artificial variabilities. These applications of morphogenic tracks in measurements of cortical thickness and cell density are important to understand the impact of gyrification on cytoarchitecture.

### Spatial profiling of multiple cell types and comparison of prefrontal cortex and secondary visual cortex

We explored the diversity in cell populations across different cortical regions by exploiting mFISH3D. We prepared three pieces of the prefrontal cortex and two pieces of the secondary visual cortex from the same donor. We labeled excitatory neurons with *Slc17a7* and oligodendrocytes with *Plp1* along with nuclear staining and α-SMA immunostaining (**Supplementary Video 5**). We detected excitatory neurons, oligodendrocytes, and all nuclei in the tissue volumes and reconstructed morphogenic tracks as we described above (**Figure 6A**). We first surveyed the differences in thickness between the two regions. We performed PCA against laminar thickness and observed the separation of morphogenic tracks between the prefrontal cortex and the secondary visual cortex (**Figure 6B**). This indicated the existence of differences in cortical thickness. Because thickness is impacted by gyrification, it is important to clarify if the differences in thickness reflected actual regional differences or arose from gyrification. Since the thickness in a gyrus/sulcus is impacted by the flux, we can minimize the influence of gyrification by taking flux into consideration. Instead of using flux, we used local flux because flux includes information on thickness in itself. We applied multivariable regression models to investigate the impact of choosing different anatomical regions on the thickness while controlling for the impact of the local flux. We used the local flux at layer IV, and found higher Cohen’s f^2^, which is the effect size in a multivariable regression model, in the deep cortical layers (**Figure 6C**). We note that Cohen’s f^2^ of more than 0.35 is generally considered large (Selya et al., 2012). This suggests larger differences occur in thickness in deeper layers than in superficial layers.

**Figure 6.**
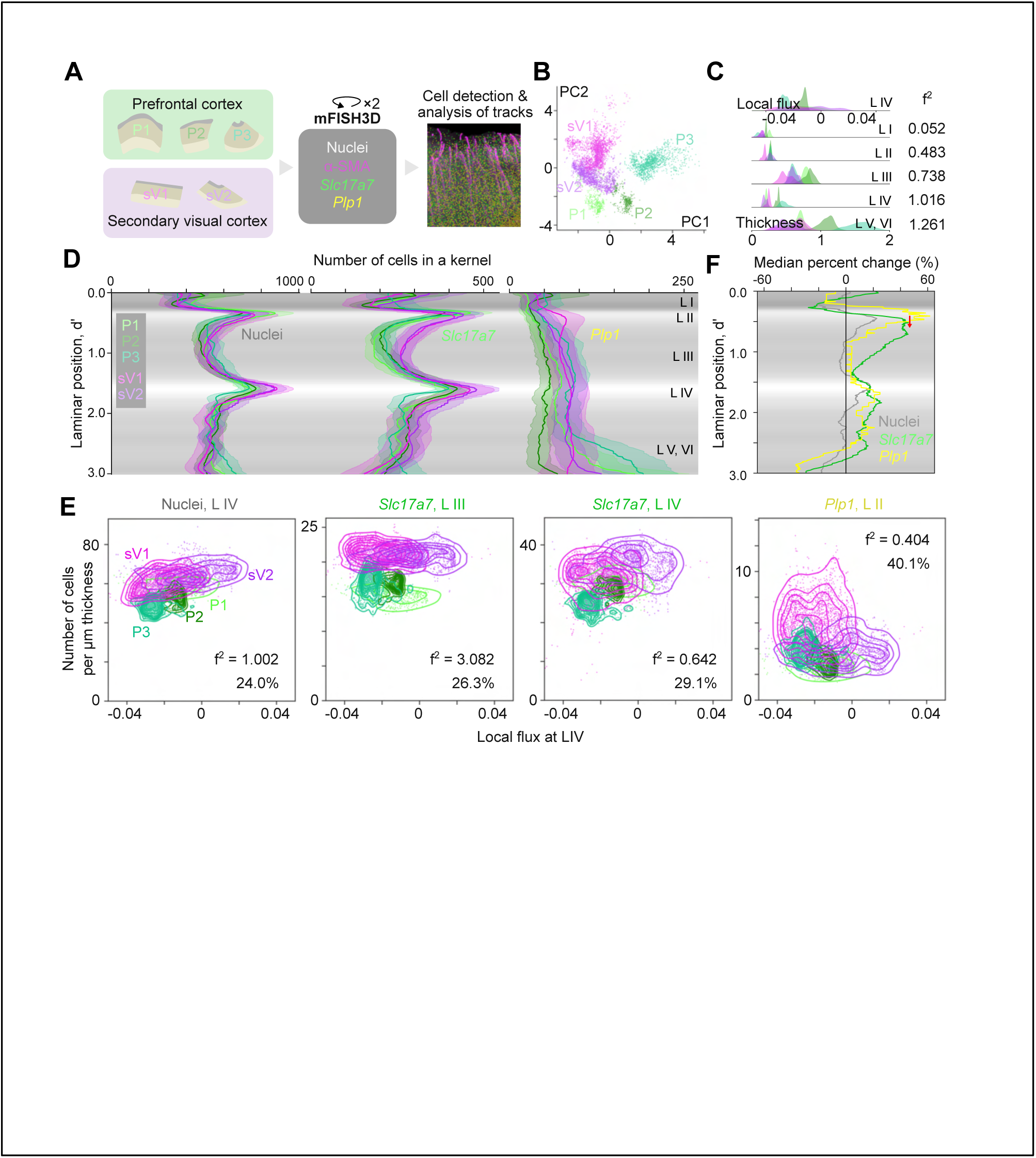
Systematic cell-type-specific density comparison between the prefrontal cortex and the secondary visual cortex. (**A**) The design of the experiment. The three pieces of the prefrontal cortex and the two pieces of the secondary visual cortex were obtained from one donor. P indicates prefrontal cortex and sV indicates secondary visual cortex. (**B**) The first and second principle components of the laminar thickness are shown for each morphogenic track. (**C**) The probability histograms of the local flux (top) and the thickness in each layer (bottom). Cohen’s f^2^ is shown on the right. (**D**) Density profiles of nuclei, *Slc17a7*+ cells, and *Plp1*+ cells by tissue pieces. The values are mean ± standard deviation (s.d.). The averaged cell density with a gray color code is shown in the background. (**E**) Scatter plot showing the local flux at layer IV and the cell densities. The plots with high Cohen’s f^2^ (more than 0.4) are shown. The median percent changes are shown along with Cohen’s f^2^. The KDE plots were overlaid on the scatter plots. (**F**) Median percent changes by cell types. The red arrow indicates the phase shift between the profiles of *Plp1*+ cells and *Slc17a7*+ cells. The averaged cell density with a gray color code is shown in the background. See **Supplementary Methods** for the calculation of median percent change. L: layer.

We investigated the difference in cell densities between the prefrontal cortex and the secondary visual cortex. We generated averaged profilings of the cell density by cell type (**Figure 6D**). We examined in which laminar positions cell density differed between the prefrontal and the secondary visual cortex by training partial least square-discriminant analysis (PLS-DA) models to predict a cortical region based on the profiles of cell densities and calculated the VIP score. From the VIP scores, we demonstrate the importance of layers II, III, and IV in the discrimination of the two regions (**Figure S6A**). To take the impact of local flux into account, we applied multivariable regression models to investigate the impact of choosing different anatomical regions on cell density, and measured Cohen’s f^2^ (**Figure S6B**). From the VIP score and Cohen’s f^2^, we expected that the cell densities were distinct in nuclei of layer IV, *Slc17a7*+ cells of layer III, *Slc17a7*+ cells of layer IV, and *Plp1*+ cells of layer II. We generated 2D plots of the densities and local fluxes to confirm the distinction and calculated Cohen’s f^2^ and median percent changes using the cell density in the layer (**Figure 6E**). We found Cohen’s f^2^ to be 1.002, 3.082, 0.642, and 0.404 for each and the transitions of the median cell densities were 24.0% 26.3%, 29.1%, and 40.1% for each. Thus our analytic scheme can detect ∼25% of regional differences in cell density. It is important to note that the profiles of the regional differences in the densities of *Slc17a7*+ cells and *Plp1*+ cells showed synchronous peaks with a slight phase shift (**Figures S6A and S6B**). The median percent change indicated that the peaks of the profile of *Plp1*+ cells appeared ∼0.2 in the laminar position (corresponding to approximately 0.2 mm in the physical distance) prior to the peaks of the profile of *Slc17a7*+ cells (**Figure 6F**). To minimize the confounding of gyrification, we grouped the morphogenic tracks with the same range of local flux and compared the tracks between the primary cortex and the secondary visual cortex (**Figures S6C**). The analysis revealed a peak shift of 0.22 in the laminar position on average. For comparison, we performed the peak-shift analysis between the profiles of the median percent change of *Slc17a7*+ cells and nuclei (**Figures S6D**). The shift was smaller and was 0.07 on average. The minor peak shift between *Slc17a7*+ cells and nuclei may have been caused by the inclusion of oligodendrocytes in the nuclei population. The major peak shift between the profiles of the median percent change of *Plp1*+ cells and *Slc17a7*+ can be explained by the myelination of the excitatory neurons. It is known that the pyramidal neurons form axon projection toward the upper layers. The higher density of excitatory neurons in the secondary visual cortex may contribute to the higher demands in myelination of the axons, and results in the higher density of oligodendrocytes at the positions slightly apart from the soma of excitatory neurons. This finding is interesting given the different developmental trajectories of excitatory neurons and oligodendrocytes (Silva et al., 2019). This synchronic emergence of *Plp1*+ cells and *Slc17a7*+ cells implies the existence of the driving factor to coordinate the population size and distribution of the oligodendrocytes in accord with the excitatory neurons or vice versa. The combination of mFISH3D and morphogenic tracks thus enabled objective regional comparison of cell ecosystems and helped to analyze cellular interactions while minimizing the confusion caused by the convoluted anatomy of a human brain.

## DISCUSSION

mFISH3D enabled scalable multiplexed whole-mount staining of a mouse brain and a human cortex. The staining protocol is based on passive immersion and easily available chemicals, allowing its widespread use for studies of three-dimensional brain anatomy. Though not demonstrated in this study, we believe the application of mFISH3D to other soft tissues such as a kidney, heart, spleen, etc., should be relatively straightforward. There are technical considerations that are important for success of mFISH3D. First, methanol pre-treatment is known to change the antigenicity of some peptides, leading to the loss of signal in immunostaining with some antibodies. This is also the case for the iDISCO immunostaining protocol, which also involves the methanol pre-treatment. There is a helpful resource list of antibodies that are compatible with iDISCO (https://idisco.info/validated-antibodies/) that may help with the choice of antibodies if immunostaining is needed. Second, the mFISH3D protocol does not support the retention of an endogenous fluorescent protein. Quenching of the endogenous fluorescent protein is thought to be due to methanol pre-treatment. Those can be alleviated by performing delipidation and RI matching in aqueous solutions (Tainaka et al., 2018, 2016). We intentionally excluded chemical modifications of the tissue to keep mFISH3D simple, but adding such chemical substances is promising for the protection of fluorescent protein and longer retention of mRNAs. Especially, the anchoring of mRNAs to the embedded hydrogel matrix is reported to enhance the durability of mRNAs, and this strategy is relatively easily integrated into our mFISH3D (Chen et al., 2016; Ku et al., 2020; Sylwestrak et al., 2016). The combination of mFISH3D and fluorescent protein is useful to reveal the relationship between projection and molecular identity of neurons (Cai et al., 2013; Livet et al., 2007). We believe the simple design of mFISH3D provides the investigators with the flexibility to customize the protocol to fit their own imaging purpose.

We proposed the concept of morphogenic tracks to evaluate the distribution of cells using the cerebral cortices of a human brain. We expect that our workflow is not limited to the cerebral cortex as far as we can extract morphogenic flows from tissues. Possible applications include cerebellum morphogenesis, spinal cord morphogenesis, and organoid morphogenesis. For example, by reconstructing the morphogenic tracks along radial glial cells in a cerebral organoid, we can recapture the migration rates or proliferation rates for radial glia from a highly heterogeneous structure of an organoid.

We demonstrated two applications of the morphogenic tracks in this study. One is to analyze the relationship between gyrification and heterogeneity of cortical thickness or cell density. So far a number of hypotheses about gyrification have been proposed mainly through macro-scale observation, but there lacks a cellular-level understanding of the mechanism. The theory that can explain every aspect of gyrification is yet to be proposed (Ronan and Fletcher, 2015). While our analysis cannot determine if the heterogenous distribution of the cortical thickness and cell density is a cause or an effect of the gyrification, analyzing the various developmental stages of brains with our workflow will bring a clearer perspective of the cellular mechanism of cortical gyrification. Cell-type specific labeling with mFISH3D will boost the understanding of the migration history of subtypes of neurons and glial cells and their impacts on gyrification. We believe that this integrated imaging scheme will help resolve the discrepancies between the theories and the cellular-level observations.

We also demonstrated the comparison of the cortical thickness and the cell populations between two functionally distinct cortical regions. We can potentially generalize the analysis further to make the analysis more consistent with biological observation (**Figure 7**). For example, we can reconstruct morphogenic tracks in the same position as main arteries or radial glial cells and can define the morphogenic tracks that adaptively change their width based on the surroundings. The comparison is not limited to cell densities. We can compare the metrics such as radial distribution function, size of cells, polarities of cells, and so on. The variety of the measurable metrics will be boosted by using modern light-sheet microscopy with isotropic and high resolution (Chhetri et al., 2015; Dean et al., 2015; Glaser et al., 2022; Kumar et al., 2014; Tomer et al., 2012; Voigt et al., 2019; Wu et al., 2013). Such analysis will help in-depth quantification of how cancer metastasis, brain stroke, and injuries affect the proliferation, cell death, and gene expression. Our workflow is expandable for the comparative analysis of multiple individuals. This is especially important to study selective cell vulnerabilities in response to neurological disorders.

**Figure 7.**
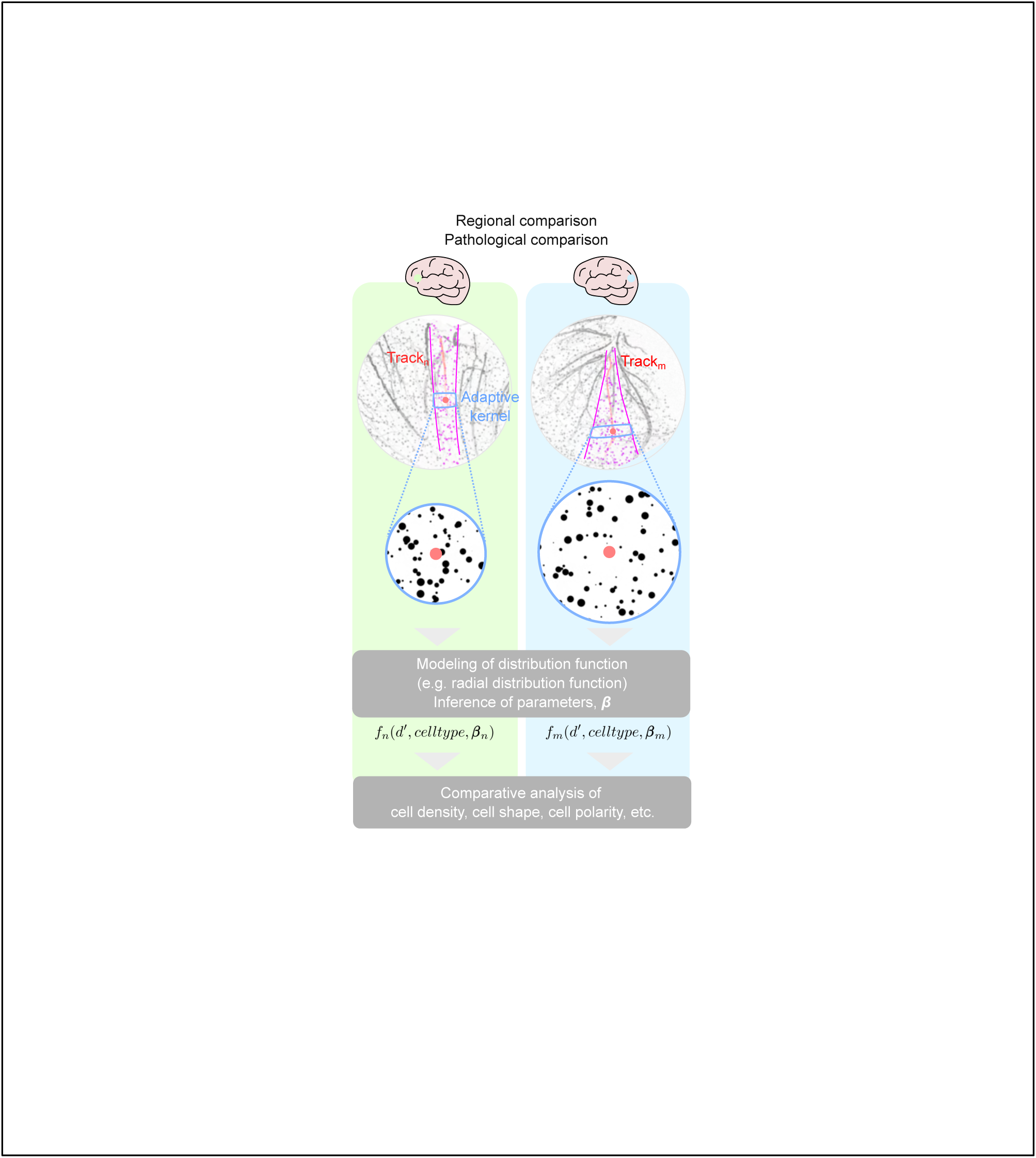
The future perspective of the comparative analysis of human brains. The researchers can place the morphogenic tracks based on the observation of the fibers, such as arteries and radial glial cells. Instead of the fixed-width morphogenic track, we can define the morphogenic track that can change the width based on the surroundings. For example, the design of the morphogenic tracks so that the cells can be allocated to their nearest radial glial cells. The modeling of the cell distribution such as radial distribution function has the potential for a detailed analysis of the impact of injuries or diseases. The quantification is not limited to cell density but can be adapted to include cell shapes and cell polarities.

Those two directions of the research, that is research in gyrification and research in comparative analysis of brains, are complementary and synergistic. Once we can predict the impact of gyrification on cell distribution, we can precisely extract genuine differences in the comparative analysis without being confounded by the gyrification. In turn, cell-type-specific developmental or evolutional comparisons of gyrencephalic brains will provide new insights to elaborate on the gyrification theory. The ability to deconvolve human brain anatomy using these methods will allow three-dimensional dissection of neurodevelopment, neurological diseases, and evolution of brains with cell-type specificity and significantly advance our understanding of these processes at the cellular level.

## ACKNOWLEDGEMENTS

This work was supported by the Howard Hughes Medical Institute (to N.H.), CHDI Foundation (to N.H.), Japanese Society for Promotion of Science Overseas Research Fellowship (to T.C.M), and Leon Levy Scholarships in Neuroscience (to T.C.M.). We thank Christina Pressl, Katherine Kane, Laura Kuss, and Maytal Babajanian for their technical assistance in the experiment on human tissues. David Davis provided us with instructions to archive human brains. Eric Schmidt proofread the manuscript. The mice tissues were prepared by Xing Jie. We thank Balakanagaram Jayaraman, Jason Banfelder, Nicholas Didkovsky, and Rebecca Bennett for their support in setting up the infrastructure of the data storage. We consulted light-sheet imaging and the downstream analysis with Alison North, Christina Pyrgaki, Katarzyna Cialowicz, Tao Tong, and Ved Sharma. We thank Kazuki Tainaka for advice on tissue clearing, Hiroki R. Ueda for advice on the chemistry of in situ hybridization, and Tomoyuki Mano for advice on configuring the workstation.

## AUTHOR CONTRIBUTIONS

T.C.M. conceptualize and designed the study. T.C.M. performed the experiments. T.C.M. analyzed the data. T.C.M and N.H. wrote the manuscript. All authors discussed the results and commented on the manuscript. N.H. acquired research funding.

## DECLARATION OF INTERESTS

The authors declare no competing interests.

**Supplementary Video 1. Whole-brain fluorescent in situ hybridization with various fluorescent molecules**

**Supplementary Video 2. mFISH3D against inhibitory neurons and oligodendrocytes with a whole-mouse brain**

**Supplementary Video 3. Reconstruction of morphogenic tracks in a human brain cortex**

**Supplementary Video 4. Segmentation of nuclei in a human brain**

**Supplementary Video 5. mFISH3D in a human brain cortex**

## MATERIALS AND METHODS

### KEY RESOURCES TABLE

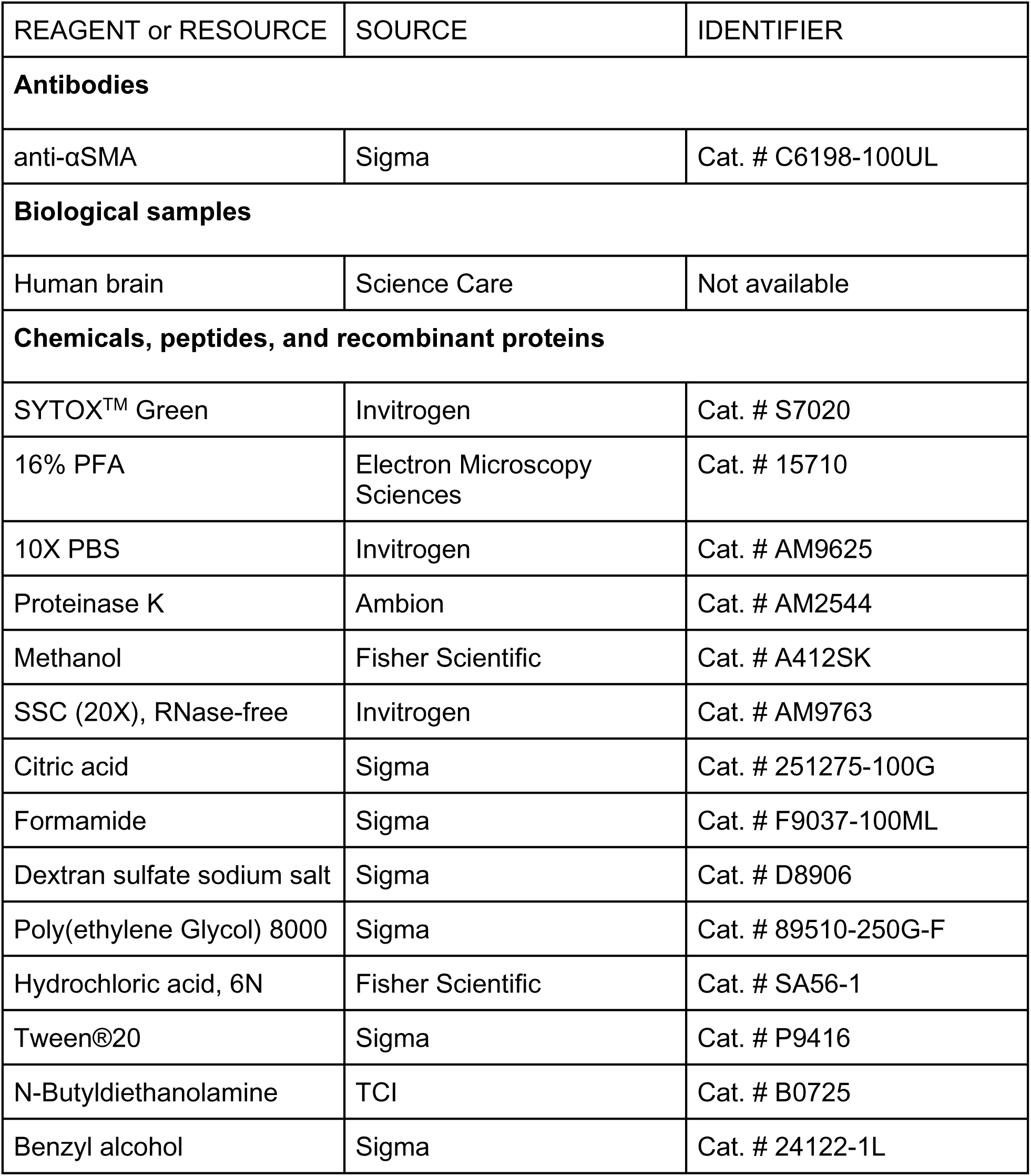
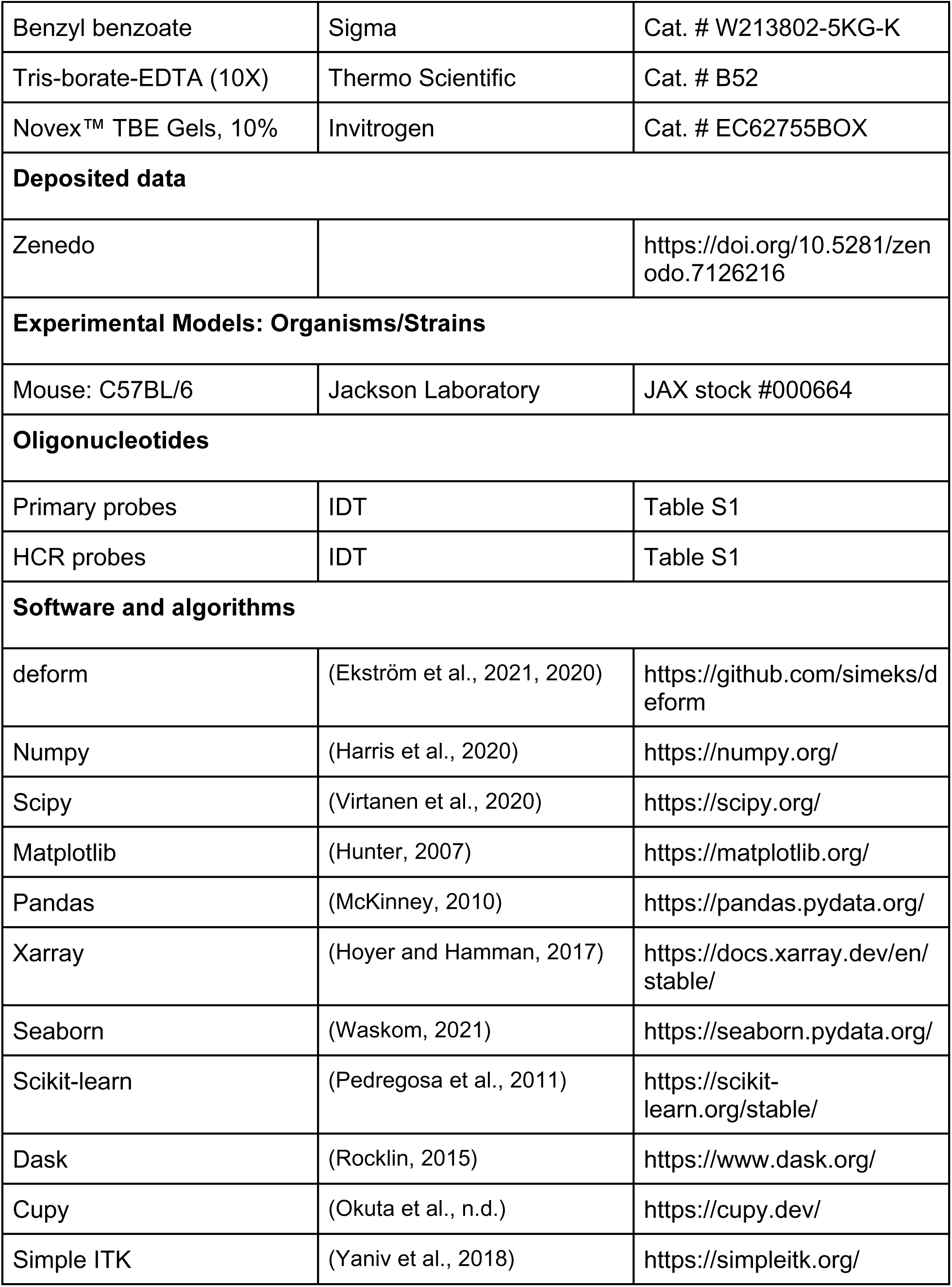
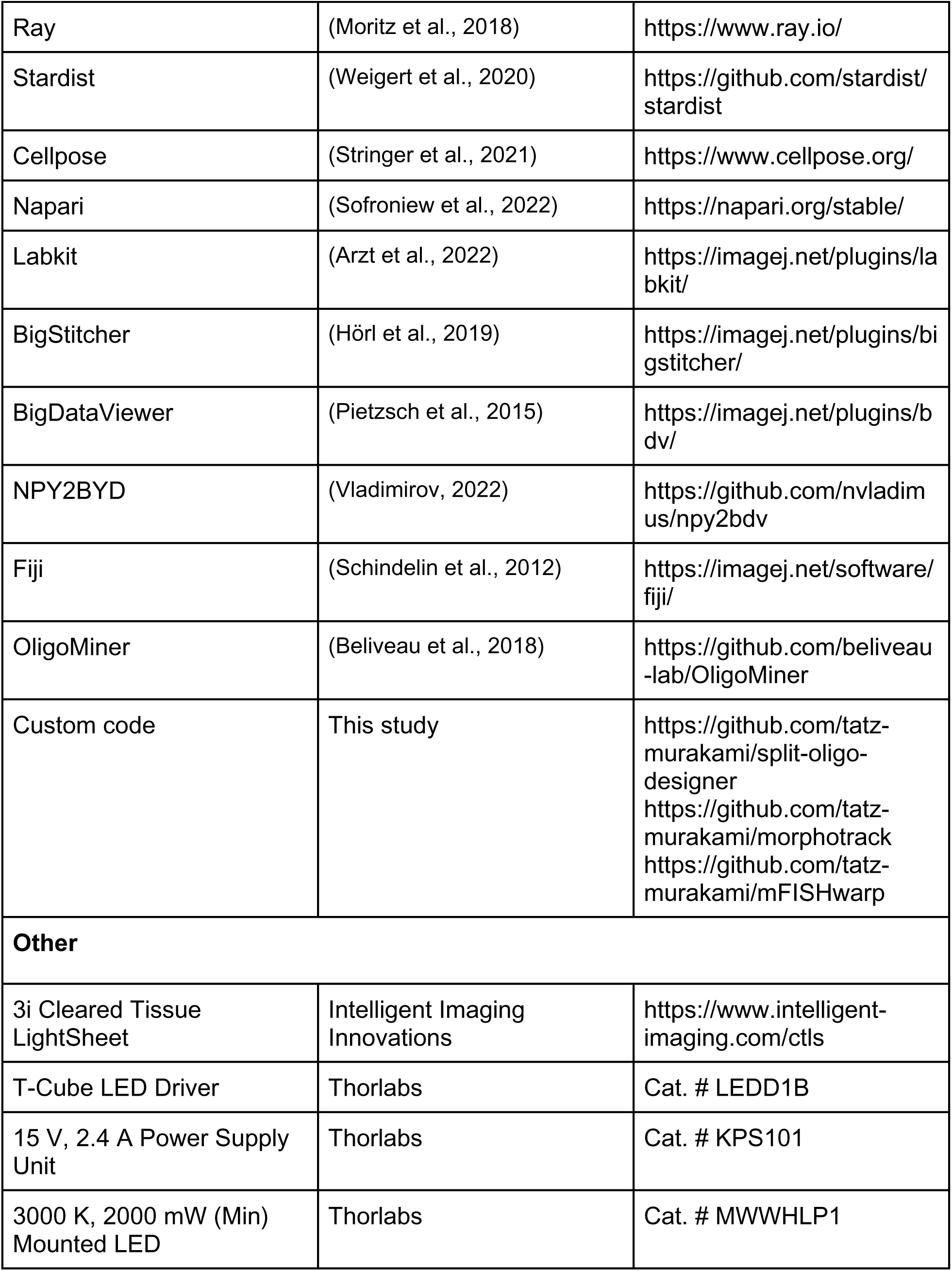

### LEAD CONTACT AND MATERIALS AVAILABILITY

Further information and requests for resources and reagents should be directed, and will be fulfilled by the Lead Contact, Tatsuya C. Murakami.

### DATA AND SOFTWARE AVAILABILITY

The transformed CUBIC-Atlas and CNN models are available at Zenodo (https://doi.org/10.5281/zenodo.7126216). Custom codes generated for this study are available on GitHub (https://github.com/tatz-murakami/split-oligo-designer, https://github.com/tatz-murakami/morphotrack, https://github.com/tatz-murakami/mFISHwarp). Due to the large size of the imaging datasets generated in the study, the datasets are not available in a public repository and are available from the authors upon request.

### MODEL SYSTEMS AND PERMISSIONS

#### Mice Model

C57BL/6J mice from Jackson laboratory were used. The mice were sacrificed by an overdose of pentobarbital (> 100 mg/kg), then transcardially perfused with 15 ml of phosphate buffer saline (PBS) and 20 ml of 4% paraformaldehyde (PFA, Electron Microscopy) in PBS. The pH of PFA-PBS was adjusted to be ∼6.5 by adding hydrochloride. The brains were dissected, then post-fixed with 4% PFA-PBS at 4°C overnight. The fixed brains were then washed with 2× saline-sodium citrate (SSC) buffer prior to the experiments. All procedures involving mice were approved by The Rockefeller University Institutional Animal Care and Use Committee (IACUC) and were in accordance with the National Institutes of Health guidelines.

#### Human Brain

A female donor 65 years of age with no known history of neuropsychiatric or neurological conditions was used in this study. The tissue samples were obtained from Science Care. The tissues were collected and banked in a fresh frozen state in accordance with approval from the local institutional review board. The tissues were provided to Rockefeller University under the authority of the institutional review board. The tissues were de-identified before receipt by Rockefeller University.

### SUPPLEMENTARY METHODS

#### mFISH3D

The most updated version of the mFISH3D protocol can be found at dx.doi.org/10.17504/protocols.io.kqdg3pjxql25/v1.

#### Design of oligonucleotides

We used in situ hybridization with signal amplification by hybridization chain reaction (Choi et al., 2018). We designed custom-made initiator-tagged DNA oligonucleotides (hereafter primary probe) by referring to the design of third-generation HCR. The design of the primary probe was modified to fit our purpose as described in the next. The sequence of mRNA was obtained from NCBI website (https://www.ncbi.nlm.nih.gov). The reverse complement of the sequence was generated. The sequence was subdivided into 20-mer oligonucleotides using OligoMiner (Beliveau et al., 2018) so that each oligonucleotide satisfies the condition: at least 1-mer interval between neighboring oligonucleotides, GC percent between 25% and 75%. We ignored the Tm value of the oligonucleotides because the split design of the primary probe can minimize the off-target signal amplification (Wang et al., 2012). To check the off-target binding, the oligonucleotides were aligned to the transcriptome database of species of interest. The database was downloaded from Ensemble (http://useast.ensembl.org/info/data/ftp/index.html). We used BLAST for the alignment. To minimize off-target signal amplification, the oligonucleotides with more than 14-mer off-target binding were subjected to further analysis. If any combination of the two off-target-binding sites from different oligonucleotides is in the proximate area, we excluded these oligonucleotides from the candidates. We set 100 bases as criteria to judge if the two oligonucleotides are proximate. The left oligonucleotides were tagged with the sequence for HCR amplification. As for the stock solution of the primary probes, we mixed 1 mM of each oligonucleotide into one Eppendorf tube. We labeled the concentration of the mixture as 1 mM primary probes for convenience. All the primary probes were synthesized by Integrated DNA Technologies (IDT). The fluorescent-dye-conjugated HCR probes were either purchased from Molecular Instruments or synthesized by IDT (Wang et al., 2020). The scripts for the design of the primary probes can be downloaded from GitHub (https://github.com/tatz-murakami/split-oligo-designer). we also designed oligonucleotides without reaction sequences of HCR (hereafter blocker probes) to form RNA:DNA hybrids. We switched the blocker probes to the reactive ones before the signal amplification. The aim of adding blocker probes is to protect mRNA from degradation while avoiding signal cross-talks.

#### Tissue preprocessing

The mouse brains were PFA-fixed by gently shaking the tissue in 4% PFA (pH 6.5∼7.0) overnight at 4°C. The human tissues were chunked into 1∼2 cm^3^ pieces of tissue blocks in the frozen state. After defrosting the tissue on ice for 30 min, the tissues were immersed in PFA-PBS (pH 6.5), then post-fixed at 4°C overnight. The tissues were sub-dissected if necessary. The fixed brains were washed with 2× saline sodium citrate (SSC) twice. The samples were treated in an increasing gradient of MeOH with gentle shaking. The concentration and durations are 30% for 30 min, 50% for 30 min, 80% for 30 min, and 100% for 30 min twice. After dehydration, the tissues were gently shaken after moving to 37°C for 2 days for delipidation. After brief rinsing with fresh MeOH, the tissues were subjected to photobleaching after clearing with the 1:2 mixture of benzyl alcohol and benzyl benzoate (BABB). The pH of BABB is adjusted to alkali by adding 1% of N-butyldiethanolamine to enhance the signal of pH-dependent fluorophores (Foster et al., 2019). We hereafter call the solvent BABB for simplicity. The cleared tissues in BABB were put under the white color LED (MWWHLP1, Thorlabs) and illuminated at full power for more than two days. We skipped this photobleaching process for mouse tissues. The photobleaching of autofluorescence was confirmed with confocal microscopy. The tissues were washed with 100% MeOH for 1 hour three times. If necessary, the tissues were stored at -30°C or -80°C.

#### Whole-mount in situ hybridization

The dehydrated tissues were re-hydrated with a decreasing gradient of MeOH (80% for 30 min, 50% for 30 min, and 30% for 30 min). The tissues were moved to 10 mM HCl and incubated for 30 min. After refreshing 10 mM HCl, the tissues were moved to 37°C and gently shaken overnight. After 30 min of washing the tissues with washing buffer (5×SSC, 0.01% Tween-20, and 20 mM citric acid, hereafter w.b.1), the solution was replaced with a second washing buffer (2×SSC, 0.01% Tween-20, and 4 mM citric acid, hereafter w.b.2) with 10 μg/ml proteinase K (AM2544, Ambion). The tissue was subjected to the gentle shake at RT for 2 ∼ 5 hours depending on the tissue size. After 30 min of washing the tissues with w.b.1, the solution is replaced with hybridization buffer (mouse brain. 5×SSC, 0.01% Tween-20, 30% formamide, and 3% PEG8000; human brain. 5×SSC, 0.01% Tween-20, and 35% formamide) for pre-hybridization. After 30-min incubation, the solution was replaced with the same hybridization buffer with 1 μM final concentration of total primary probes. The tissues were moved to 37°C and gently shaken overnight or for 2 days depending on the tissue size. The tissues were washed with w.b.1. If we see a non-specific signal in a human brain, we added 10% formamide to w.b.1. After the 2 hours of washing three times, we replaced the solution with the hybridization buffer for pre-hybridization. After 30 min, the buffer was replaced with the hybridization buffer with 90 nM final concentration of HCR probes. The HCR probes were denatured at 95°C for 90 seconds and snap-cooled at RT for 30 min beforehand. The amplification was performed overnight or for 2 days depending on the tissue size at room temperature. The buffer was replaced with w.b.1 and the tissue was washed for 1 hour three times. For human tissues, we added SytoX^TM^ Green and 1/1000 concentration of anti-αSMA antibody to w.b.1 and incubated for 2 days at room temperature. The w.b.1 was replaced with w.b.2 with 1-hour incubation. After treating the tissues with the increasing gradient of MeOH (30% for 30 min, 50% for 30 min, 80% for 30 min, and 100% for 30 min twice), we immersed the tissue in BABB until the tissue became fully cleared.

#### Multi-round whole-mount in situ hybridization

To minimize the breeding of the residual signals from the last round, we optionally included photobleaching after the first round of imaging. The photobleaching was performed overnight. The tissue was de-cleared by 100% MeOH and treated with a decreasing gradient of MeOH as stated above. After 30% MeOH, the tissue was washed with w.b.1 for 30 min twice then treated with hybridization buffer for 30 min for pre-hybridization. The buffer was replaced with 1 μM final concentration of total primary probes for the second round in the hybridization buffer. If there can be possible cross-talk between the primary probes of the previous round and the HCR amplification of this round, we performed heat-assisted de-hybridization to remove the primary probes of the previous round. The tissue was heated to 65°C for 1 hour. The buffer was then refreshed again with the hybridization buffer with the primary probes. We skipped this de-hybridization step if there are no concerns about the cross-talk. The second-round hybridization was performed at 37°C overnight or for 2 days depending on the tissue size. We follow the same procedures for the amplification and clearing as stated above. If needed, the process is repeated.

#### Whole-mount immunostaining

The tissues were re-hydrated with a decreasing gradient of MeOH. The concentration and durations are 80% for 30 min, 50% for 30 min, and 30% for 30 min. The tissues were moved to w.b.1 and incubated for 30 min. After refreshing w.b.1, the antibodies or nucleus dyes were added. In this study, 1/1000 diluted Cy3-conjugated anti-αSMA (C6198-100UL, Sigma Aldrich), and 1/5000 diluted SytoX Green (S7020, Thermo Fisher Scientific) were used. The staining was performed at room temperature (RT, 20 ∼ 25°C) with a gentle shake overnight or for up to 2∼3 days depending on the size of the tissue. The samples were cleared with BABB after dehydration and subjected to imaging.

#### Staining strategy

The combination of the dyes and oligonucleotides, and strategies are described in **Table S1**.

#### Light-sheet microscopy

The Cleared Tissue LightSheet (CTLS) system from Intelligent Imaging Innovations was used for all the imaging of cleared tissues. The system is equipped with fiber-coupled lasers (405, 488, 561, 640, and 785 nm). A spatial light modulator is used to modulate the beam shape and the light sheet is created by a galvanized mirror. The illumination objectives 5×/0.14NA are on the left and right sides of the imaging chamber. The imaging chamber is filled with BABB (refractive index = 1.56). The image is captured through an objective lens (Zeiss, PlanNeoFluar Z 1×/0.25NA FWD=56 mm). To enable the high-speed transition of multiple illumination wavelengths in one plane, the system is equipped with a high-speed filter wheel (OPTOSPIN). ORCA-fusion BT is used as an sCMOS sensor.

We switched the illumination wavelength at every z position and the chromatic focus shift is dynamically adjusted by the spatial light modulator. The stack-wise z-scan was performed and the motorized stage repositioned the sample in XY-direction to cover the whole tissue volume. We used static illumination focus during the imaging. We added 5× or 10× optical zoom (5× for mouse tissues and 10× for human tissues) ending up 1.3 μm or 0.65 μm pixel size. The 3 μm z-step was chosen. All the hardware was controlled by Slidebook (Intelligent Imaging Innovations). The data was stored at PowerEdge R740XD (Dell) with twelve 16 TB HDDs through 10-GB Ethernet fiber.

#### Data analysis system

We used a DGX A100 with four NVIDIA A100 40GB GPUs for the image analysis.

#### Stitching and correction of chromatic aberration

The CTLS produced z-stacks in Numpy array binary file format (.npy). We converted the .npy files to hierarchical data format (.h5) with metadata of stage positions (.xml) using NPY2BDV (https://github.com/nvladimus/npy2bdv). The resulting .h5 contains multi-resolution pyramids of the images. We used BigStitcher Fiji plugin for the stitching and the correction of chromatic aberration (Hörl et al., 2019). In detail, we first performed translational stitching using the channel of ribosomal RNA or nucleic staining. The rigid transformation is usually not enough to produce stitching in a single-cell precision, so we also performed affine transformation based on the iterative closest point after detecting feature points with difference of gaussian filters. We used the default parameters for the process and used the downsampled images (downsampled by 8, 8, and 4 times in x, y, and z each). Importantly, we applied the iterative closest point (ICP) stitching for the correction of tile shifts and chromatic aberration altogether. If this did not produce a satisfactory result, we corrected the chromatic aberration z-stack by z-stack with ICP and then applied a second ICP to fix tile shifts. We further refined the affine transformation by the next round ICP with the higher resolution of images (downsampled by 4, 4, and 2 times in x, y, and z each). Our visual inspection confirmed this scheme could produce both the stitching and correction of the chromatic aberration at single-cell precision in all the imaging cases. We blended the overlapped regions and exported 16-bit unsigned integer fused images. We convert the images to .zarr with multi-resolution pyramids for the following analysis.

#### Registration

The image registration of large 3D in a single-cell precision was reported previously (Wang et al., 2021). Bigstream is based on three rounds of image registration; 1) global affine registration, 2) chunk-wise affine registration, and 3) chunk-wise non-linear registration. The global registration is performed using downsampled images and the chunk-wise registration is performed by dividing the high-resolution images into chunks. The chunk-wise registration is necessary if the data size cannot fit into the system RAM and this makes the strategy appropriate for light-sheet imaging data. This strategy assumes initial global affine registration can align overall shapes to perform second chunk-wise affine registration. Otherwise, the regional loss of the images occurs. In our case, because of the tissue size, the tissue can cause mild bending around at pons of mouse brains and violate the assumption. Moreover, the non-linear registration of Bigstream does not utilize GPUs, making the process slow. Alternatively, we decided to perform 1) global affine registration, 2) global non-linear registration, and 3) chunk-wise non-linear registration with GPU acceleration (Ekström et al., 2021, 2020). The downsampled image (downsampled by 8, 8, and 8 times in x, y, and z) was used for both 1) and 2). The dense intensity-based affine registration was performed with the cross-correlation as a similarity metric. We used the affine transformation matrix for the initial alignment of the global non-linear registration with the cross-correlation as a similarity metric. The obtained displacement field was upsampled for the following chunk-wise registration. We used the higher resolution image (downsampled by 2, 2, and 2 times in x, y, and z) for the chunk-wise non-linear registration. Though we could perform the registration at the original resolution, we did not observe improvements in the quality of the registration. The images were subdivided into chunks with 32 voxels of overlaps. We used the upsampled displacement fields from the previous step as the initial alignment and performed chunk-wise registration. We used cross-correlation as a similarity metric. The obtained chunks of displacement fields were linearly fused and were used to produce the final warped image. The scripts were uploaded to GitHub (https://github.com/tatz-murakami/mFISHwarp).

#### Segmentation of cells and detection of centroids

##### Choice of CNN models

We used Stardist and Cellpose for the segmentation purpose of mFISH3D-stained cells. Both Stardist and Cellpose are python packages for CNN-based cell segmentations with different advantages and weaknesses. We chose Stardist for the mouse brains. Stardist can train the CNN model with three-dimensional images. Because our data has an uneven axial resolution in a single plane due to the shape of the gaussian beam, training with three-dimensional images helps to make a model which can segment the cells in an isotropic way regardless of the position of the cells. This is important to minimize the false positive of the satellite oligodendrocytes as shown in **Figure 3**. We also note that at our mesoscale resolution for mouse brains (voxel size: 1.3 μm × 1.3 μm × 3 μm), Cellpose caused over-segmentation while we did not see the issue in Stardist. The downside of the Stardist is the limited scalability of the prediction. Non-maximum suppression (NMS) is used for the prediction of the cells, but the performance of NMS becomes deteriorated according to the increase in the number of segmentable objects. The lack of GPU implementation of NMS in Stardist also slows down the prediction. For this reason, we used Cellpose in nuclei and cell segmentations of a human brain. To avoid the over-segmentation issue and for the better separation of individual nuclei, we imaged the human brain with higher magnification (voxel size: 0.65 μm × 0.65 μm × 3 μm). We did not see over-segmentation issues. Cellpose performs the segmentation according to the predicted gradient of the objects with the GPU acceleration, making the segmentation process faster than Stardist.

##### Training of models

We trained the Stardist model using 3D training datasets. Because of the distinct morphologies, we decided to make two models: one model for neurons and another model for oligodendrocytes. The training datasets were made by manually annotating several hundreds of cells. We made isotropic-sized annotations regardless of the beam waists. The F1 score of the model for inhibitory neurons was 0.82, and the F1 score of the model for the oligodendrocytes was 0.92 in the cortex. We trained the Cellpose model using the 2D training dataset because Cellpose does not require 3D training datasets for 3D segmentation. Based on the morphological distinction, we decided to make three models: one model for nuclei, one model for neurons, and another model for oligodendrocytes. The training datasets were made by manually annotating several hundreds of cells. Because the human brain produces more unspecific noises from vasculatures compared to mouse brains probably due to the incapabilities of cardinal perfusion, we needed to take additional measurements to minimize the false positives. We used the fact that the signal-to-noise ratio of nuclei staining is higher than that of mFISH3D. By including the nucleic signal in the datasets of neurons and oligodendrocytes, and by training the model with dual-channel datasets, we could reduce the number of false positives. The F1 score of the model for nuclei was 0.92, for *Slc17a7*+ cells was 0.85, and for *Plp1*+ cells was 0.92. The trained StarDist models and Cellpose models can be downloaded at Zenodo (https://doi.org/10.5281/zenodo.7126216).

##### Normalization

Due to the scattering or uneven diffusion of the fluorophores, the signal intensity is often uneven over the tissue. The normalization of the intensity is necessary to obtain consistent accuracy of the segmentation. The normalization using the local intensity is one of the popular choices for light-sheet data (Matsumoto et al., 2019).

Matsumoto et al. performed the local normalization by assuming a kernel of a certain size. The voxel-wise normalization could be performed by calculating the local minimum and local maximum within the kernel. We initially tested the approach and found some of the cells were left unsegmented. The reason is that if the true signals exist just next to the unspecific signal with high intensity, the true signal is diminished due to the local normalization and ends up in false negatives. Since it is common to have unspecific signals from autofluorescence of blood vessels or myelin, an alternative normalization was wanted. Instead of using the local maxima using a kernel, we decided to use the intensity of the neighboring cells. First, we apply segmentation using kernel-based normalization. This leaves some cells unsegmented for the reason stated above but can segment the majority of the cells without segmenting the non-cell objects. The normalization was fixed using the intensity of the neighboring segmented objects as the local maximum. The second round of segmentation was applied to the updated normalized images. We found this could largely reduce false negatives. We note that this doubles the computation time.

##### Segmentation and stitching of segmentation

The instance segmentation was performed by dividing the high-resolution images into chunks with overlaps and by applying the models to the chunks. The downside of chunk-wise segmentation is that it can produce disconnected segmentation if the cells exist across multiple chunks, and this could produce over-segmentation. To minimize the issue, we took overlaps so that at least the whole shapes of cells appear at least in one chunk, and merged segmentation using IoU as a criterion. We used the IoU threshold 0.7 and fused the segmentation. The segmented objects were renumbered so that each object has a unique identification number. The centroids of the segmented objects were then extracted using “regionprops” function in the scikit-image python library.

#### Classification of inhibitory neurons

For the analysis of **Figures 2** and **3**, we classified the inhibitory neurons based on the expression of the marker genes. Because it is known that some *Sst*+ cells are *Pvalb*+ cells and some *Pvalb*+ cells are not inhibitory neurons, we classified the cells by following the next few steps. We used the segmentation result of *Gad1*+ cells. All the *Gad1*+ cells were regarded as inhibitory neurons. During the segmentation process, Stardist produces probabilities of the objects. We measured the probabilities of being *Vip*+, *Sst*+, *Pvalb*+, and *Npy*+ cells. The classification was performed by measuring these probabilities in the segmentation of *Gad*1+ cells. We first identify the *Vip*+ cells if the probability within the segmented object was higher than the threshold. We used the threshold that was calculated during the model training of Stardist. All the *Gad1*+ cells except *Vip*+ cells were used for the classification of *Sst*+ cells. Using the same strategy as the identification of *Vip*+ cells, we identified *Sst*+ cells. By excluding *Vip*+ and *Sst*+ cells, we identified *Pvalb*+ cells again with the same strategy. We labeled the left *Gad*+ cells as “Other”. Using the probability of *Npy*+ cells within the segmentation of *Gad1*+ cells, we sub-classified the inhibitory neurons into *Npy*+ and *Npy*-.

#### Simulation of the virtual tissue sections

Using SytoX^TM^ Green nuclear staining, we performed imaging of the ∼4×4×3 mm^3^ human cerebral cortex. The nuclei were segmented using Cellpose and their centroids were detected according to the method described above. The tissue slice with 500-μm width and 50-μm thickness were simulated by changing the sectioning angle by one degree while fixing the position of the pivoting axis. We visually confirmed the reconstructed tissue sections cover layer I to white matter. The layer annotations of the centroids were obtained by measuring the layer annotations of the nearest morphogenic track. The centroids that were included in the virtual section were used for the quantification of the cell number and density.

#### Reconstruction of morphogenic tracks

The scripts for the reconstruction of morphogenic tracks are available on GitHub (https://github.com/tatz-murakami/morphotrack). For the simulation and analysis of morphogenic tracks, we used downsampled images with 10 μm × 10 μm × 10 μm voxel size and convert the coordinates of the cells into 10-μm units for consistency.

##### Construction of a vector field

The artery signals from the alpha-smooth-muscle actin antibody are used. The arteries were segmented using Labkit (Arzt et al., 2022). The segmented arteries were skeletonized. The local vectors were calculated on each voxel of the skeletonized arteries based on the averaged vectors between k nearest neighbors. The calculated local vectors were allocated to the segmented arteries by nearest neighbor interpolation so that the thicker arteries gain more weight in the following steps. While the major branches of the artery go along a morphogenic flow, other minor branches stochastically distribute from the major branches and form sub-branches. By averaging the local vectors within a certain size of a region, we could cancel out and minimize the impact of the minor branches. We used a spheric kernel with a radius of 420 μm for the purpose. After averaging, the vector fields were constructed by the polynomial fitting of each component of the vectors. We used a fifth-degree polynomial (**Figure S4D**).

##### Selection of seeds

We used the outermost cells of layer 1 as seeds. To detect the outermost cells, we generated an alpha shape from the centroids of the nuclei. The vertices on layer 1 were selected. To avoid the oversampling of the morphogenic tracks, we randomly chose 10% of the vertices and used them as seeds.

##### Reconstruction of morphogenic tracks

The morphogenic tracks were simulated by numerically solving the ordinal differential equation (ODE).

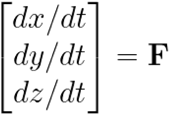

Where **F** is the reconstructed vector field. Note that **F** is independent of *t*. For each morphogenic track, we calculated local cell densities at *t*. We defined a disk-shaped kernel, which has 100 μm in radius and 20 μm in height. We count the number of cells within the kernel and used the counts as local cell density. The orientation of the kernel is determined to be consistent with the local vector of the simulated vector fields.

##### Alignments of morphogenic tracks

Note that *t* is defined for the sake of the simulation and it does not have any biological meaning. We could convert *t* to the distance from the tissue surface (*d*), but the metric is not convenient to indicate the laminar position of the cortex. The alignments are non-linear transformations that convert *t* or *d* to transformed distance (*d*’) so that the same laminar positions are comparable among morphogenic tracks. We used the local densities of nuclei on morphogenic tracks for the alignment. The alignment is composed of two steps; linear transformation followed by non-linear transformation. We linearly transformed the *t* so that the tracks are at the same position at the outermost of the tissue and the border of the gray matter and the white matter. The linearly aligned tracks were non-linearly transformed using the same algorithm we used in the image registration but in a single dimension. We chose one representative morphogenic track and aligned other tracks to the representative track. To avoid the registration from overfitting, we removed the high-frequency component using Fourier transformation. Using cosine similarity and mutual information as similarity metrics, we rejected the morphogenic tracks with poor alignment qualities. The quality of the alignment was visually inspected.

##### Surface flattening

We used the positions of the seeds for the surface flattening. Isomap with 20 neighbors with the two components was used. The surface flattening was performed for visualization purposes. The origin of the surface-based coordinate is arbitrarily chosen since it does not affect our analysis and interpretation of the data.

##### Calculation of flux and local flux

The flux is obtained from the vector fields of the morphogenic flow given a surface *S*. We normalized the strength of the vector field so that the strength is even in any position. The normalization was performed so that the size of the vector is the same. If we assume that the vector fields represent the flow of a fluid, the flux represents the amount of fluid flowing through the surface *S* per unit of time. We determined the sign of the flux so that the positive flux means the convergence of the morphogenic flows while the negative flux means the divergence of the morphogenic flows. We parameterized the *S* with a curve *C*, which is usually a section of a morphogenic track, and *r*, which is the distance from the curve *C*. We used fixed *r* (= 100 μm) throughout the study. We also defined local flux which is the derivative of flux with respect to the length of *C* (**Figure 5D**).

##### Limitations

We used polynomial curve fitting for the reconstruction of the vector field. Though the approach worked fairly well in our tissue size, the fitting may not be appropriate for the larger and more complex convolved shape. We noticed our strategy of choosing the positions of the seeds tends to cause oversampling in gyri while sparse sampling in sulci. This can be alleviated by adjusting the sampling frequency adaptively based on the local density of arteries or local flux. The alignment algorithm guarantees the differentiable displacements along a track but does not guarantee the differentiable displacements in the original XYZ space. This is because the alignments were performed track-wise, and the strategy did not consider the displacements of nearby tracks. Further sophistication of the displacement strategy is necessary when differentiable displacements are wanted.

### QUANTIFICATION AND STATISTICAL ANALYSIS

#### Statistical analysis

The statistical analysis was conducted with the Scipy Python library (Virtanen et al., 2020). The error bars indicate standard deviation throughout the manuscript.

*P values*. In **Figure 3C**, one-way ANOVA with Tukey’s post hoc test for multiple comparisons is used. N represents the number of mice used.

*Cohen’s f^2^*. Each morphogenic track was regarded as an individual sample. We made a multiple linear regression model with a label of the region and local flux as covariates to predict cortical thickness (**Figure 6C**) or cell density (**Figures 6E** **and S6B**). Cohen’s f^2^ of the label of the region was calculated from the model.

*VIP score*. See the PLS regression section below.

*Median percent change*. The tracks were grouped based on the regions (**Figures 6E** and **6F**). If necessary, the tracks were subgrouped based on the local flux (**Figures S6C and S6D**). For the subgrouping, the percentile range of the bin size of 10% was used. The medians of the cell densities at each laminar position were calculated to obtain the median density profile, *d*. The median density change was calculated according to the following formula where *d_sV_* indicates the median density profile of the secondary visual cortex and *d_p_* indicates the median density profile of the primary cortex.

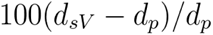

#### CUBIC-Atlas

CUBIC-Atlas version 1.2 was downloaded from http://cubic-atlas.riken.jp. Because the CUBIC-Atlas was generated from an expanded brain, the coordinates have discrepancies from other standard mouse brain atlas such as Allen Mouse Brain Atlas CCFv3. To keep the consistency in our analysis, we transformed the coordinates of CUBIC-Atlas to CCFv3 by non-linear registration. The transformed coordinates were used for the analysis. The transformed CUBIC-Atlas can be downloaded from (https://doi.org/10.5281/zenodo.7126216). Prior to the probabilistic annotation, we registered the signals of ribosomal RNA to CCFv3. The annotation and k-means clustering are performed according to the method described in the previous study (Murakami et al., 2018).

#### Theoretical probability and empirical probability

The local densities of each cell type around candidate oligodendrocytes were calculated for the theoretical probabilities. To calculate empirical probability, we need a number of candidate oligodendrocytes with the exact same theoretical probabilities. Because such incidence is uncommon, we grouped the two-hundred candidate oligodendrocytes with the nearest theoretical probabilities and calculated the empirical probability per one candidate oligodendrocyte.

#### PLS regression and PLS-DA

In **Figure 5K**, the PLS regression was used to make a model for predicting fluxes using the cell densities. For the determination of the number of the principal component to use in the prediction, we performed 10-fold cross-validation for each choice of the components. The best model with the lowest mean square error was chosen for the analysis of the VIP score. The VIP score was calculated according to the past report (Eriksson et al., 2006, p.). For the classification problem, we used the PLS-DA models (**Figure S6B**). The PLS regression is trained to predict the label of the region using the cell densities. Cross-validation was performed for the selection of the best model as stated above.

#### Peak-shift analysis

The peak shifts were quantified in **Figures S6C and S6D**. The profiles of median percent changes of *Plp1*+ or nuclei were non-linearly aligned to the profile of *Slc17a7*+ in the same way described in the alignments of the morphogenic tracks. The first major peak was detected in both the original profile and warped profile. The distance between the first major peaks was measured.

---

**Figure S1.**
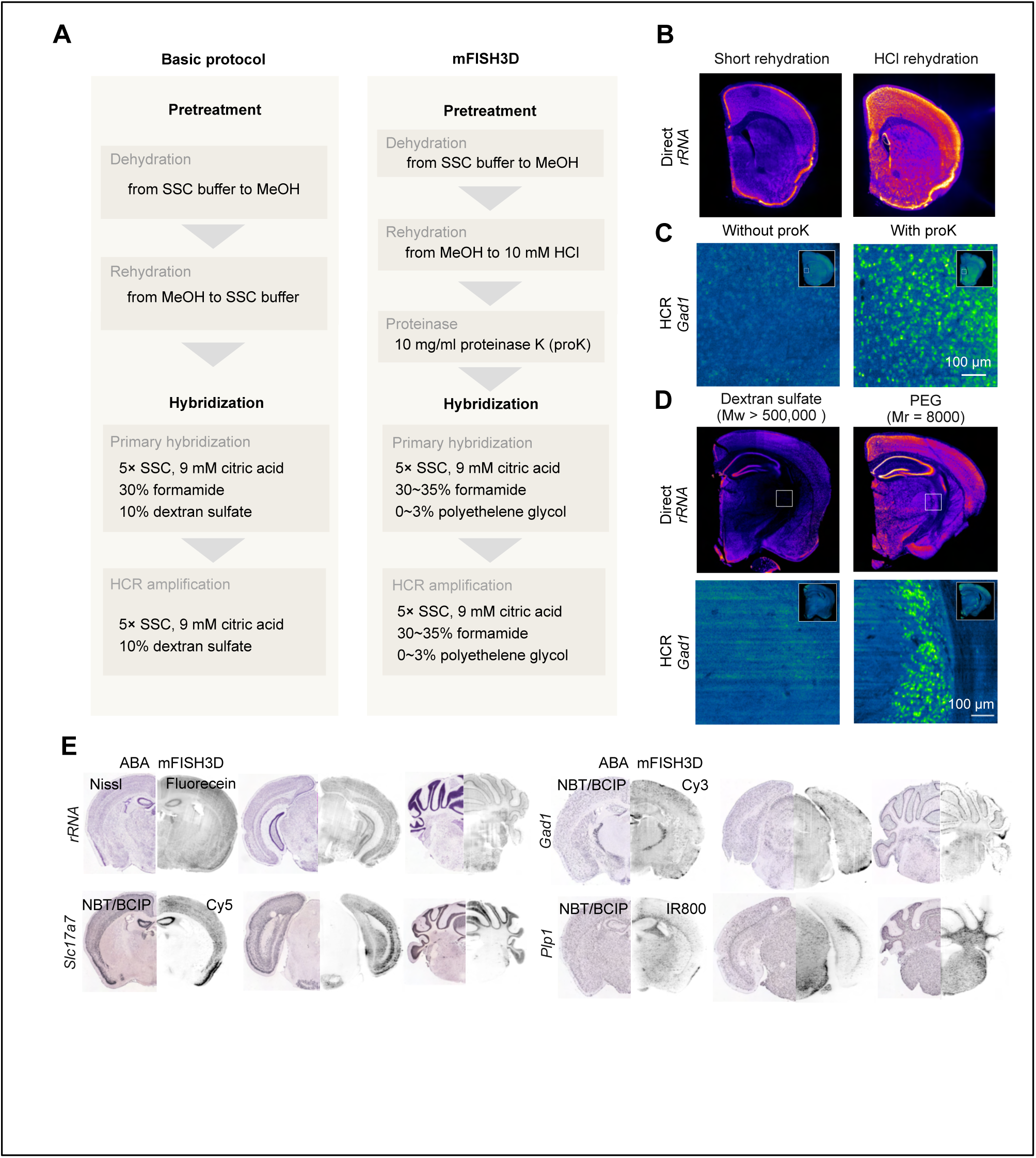
Chemical exploration of 3D in situ hybridization for enhanced signal-to-noise ratio. (**A**) The comparison of the basic protocol and mFISH3D protocol. The chemical compositions of each reaction step are shown. (**B**) The impact of mild acidic rehydration. The short rehydration shows a weak fluorescent signal inside the tissue. The coronal tissue blocks from a mouse brain with 2-mm-thickness were used. The signal intensity is visualized in pseudocolor. (**C**) Pretreatment with proteinase K prior to hybridization of primary probes. The tissue blocks from a mouse brain with 2-mm-thickness were stained by targeting mRNA of *Gad*1 and visualized in pseudocolor. The signal was amplified with HCR. The overviews of the tissues are shown in the insets. (**D**) Polyethylene glycol as an alternative to dextran sulfate in hybridization buffer. The coronal tissue blocks with 2-mm-thickness are stained by targeting *rRNA* and mRNA of *Gad*1 and visualized in pseudocolor. The signal of *Gad1* was amplified with HCR. The overviews of the tissues are shown in the insets. The optical sections at 500-μm-depth from the dissected positions are shown in **B**, **C,** and **D**. (**E**) mFISH3D against mouse hemisphere brains. The reconstructed coronal sections are shown at three different positions for each staining of *rRNA*, *Gad1*, *Slc17a7*, and *Plp1*. The screenshots of ISH data from Allen Brain Institute are shown in the left half for comparison.

**Figure S2.**
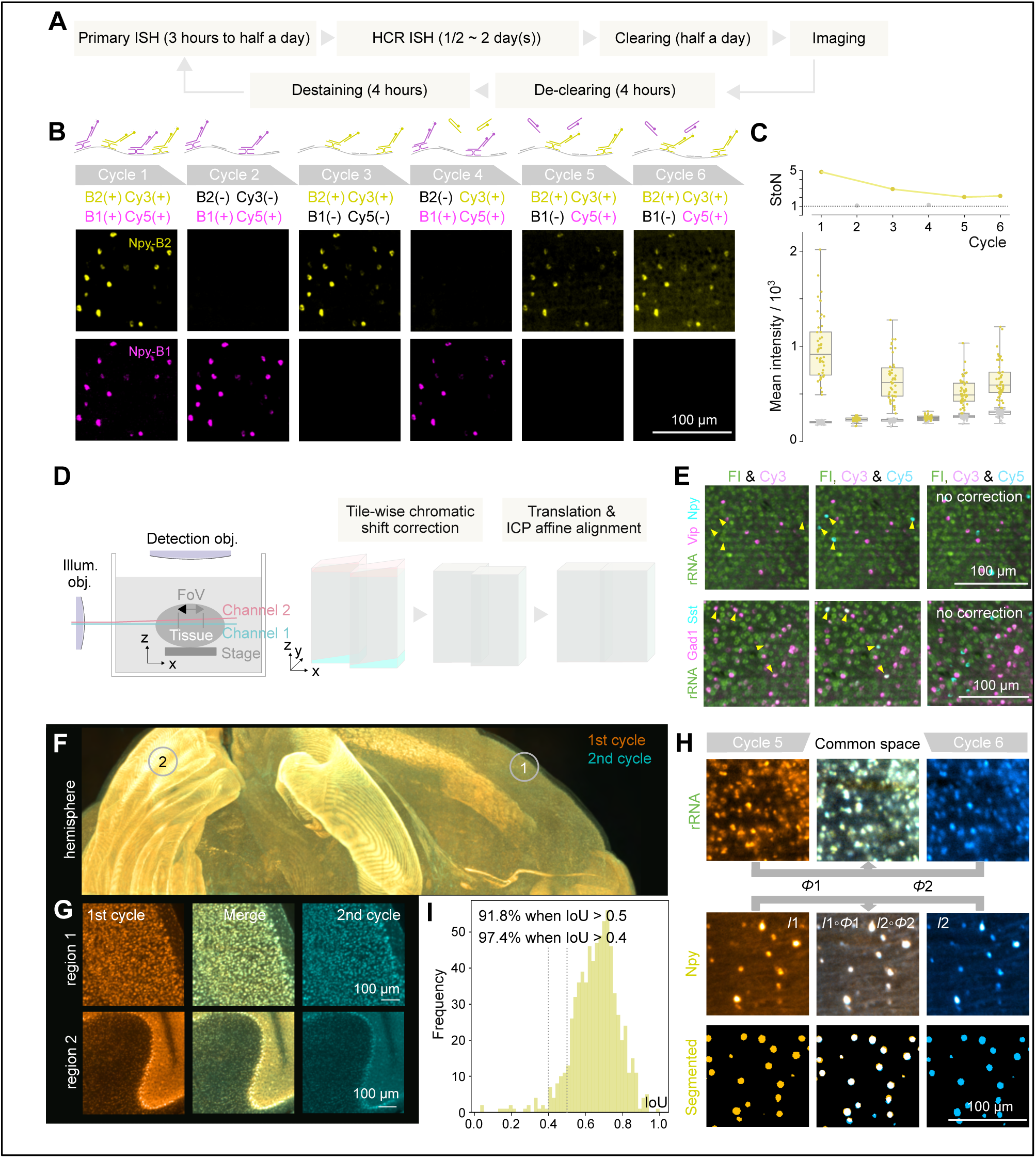
Workflow of the image analysis of mFISH3D. (**A**) The strategy to achieve multiplexed imaging of mRNAs with mFISH3D. (**B**) The carry-over of the primary probes and fluorophores from the previous cycles was examined. The data also suggested that the signal can be amplified after multiple cycles of hybridization and clearing processes. A 2-mm-thick mouse-brain coronal block was used and the magnified views are shown. (**C**) The quantification of the fluorescent intensity from the dataset of **B**. For the quantification, we made a number of ROIs that covered the cells or background. The signal-to-noise ratio (StoN ratio) was quantified by taking the ratio of the signal intensities in cells and background intensities (top). The mean intensity of each ROI was plotted (bottom). The box plot indicates minimum, lower quartile, median, upper quartile, and maximum. (**D**) Illustration of one of the major causes of the chromatic focus shift (left). Because the refractive index of clearing solvent is a function of wavelength, the illumination of different z-plane can be induced. The chromatic focus shift can be further exacerbated by the chromatic aberration of the detection lens. We fixed the chromatic focus shift for better stitching of the tiles (right). (**E**) The representative magnified views of the mouse brains with three-color imaging. The yellow arrowheads indicate cells with double/triple signals. The images without chromatic corrections are shown for comparison (right). (**F**) The cellular-resolution registration of the images from different cycles. The maximum z-projection of a mouse hemisphere brain is shown after warping the 2nd cycle to match the 1st cycle. (**G**) The magnified views of the **F**. The cerebral cortex (region 1) and the cerebellum (region 2) are shown. (**H**) The transformation field (Φ1 and Φ2) obtained through registering the *rRNA* channels from two cycles was applied to the *Npy* channel. The cells were segmented based on the signal of *Npy*, and the intersection over union (IoU) of the segmentations of Cycle 5 and Cycle 6 was examined. (**I**) The quantification of IoUs is visualized in the histogram.

**Figure S3.**
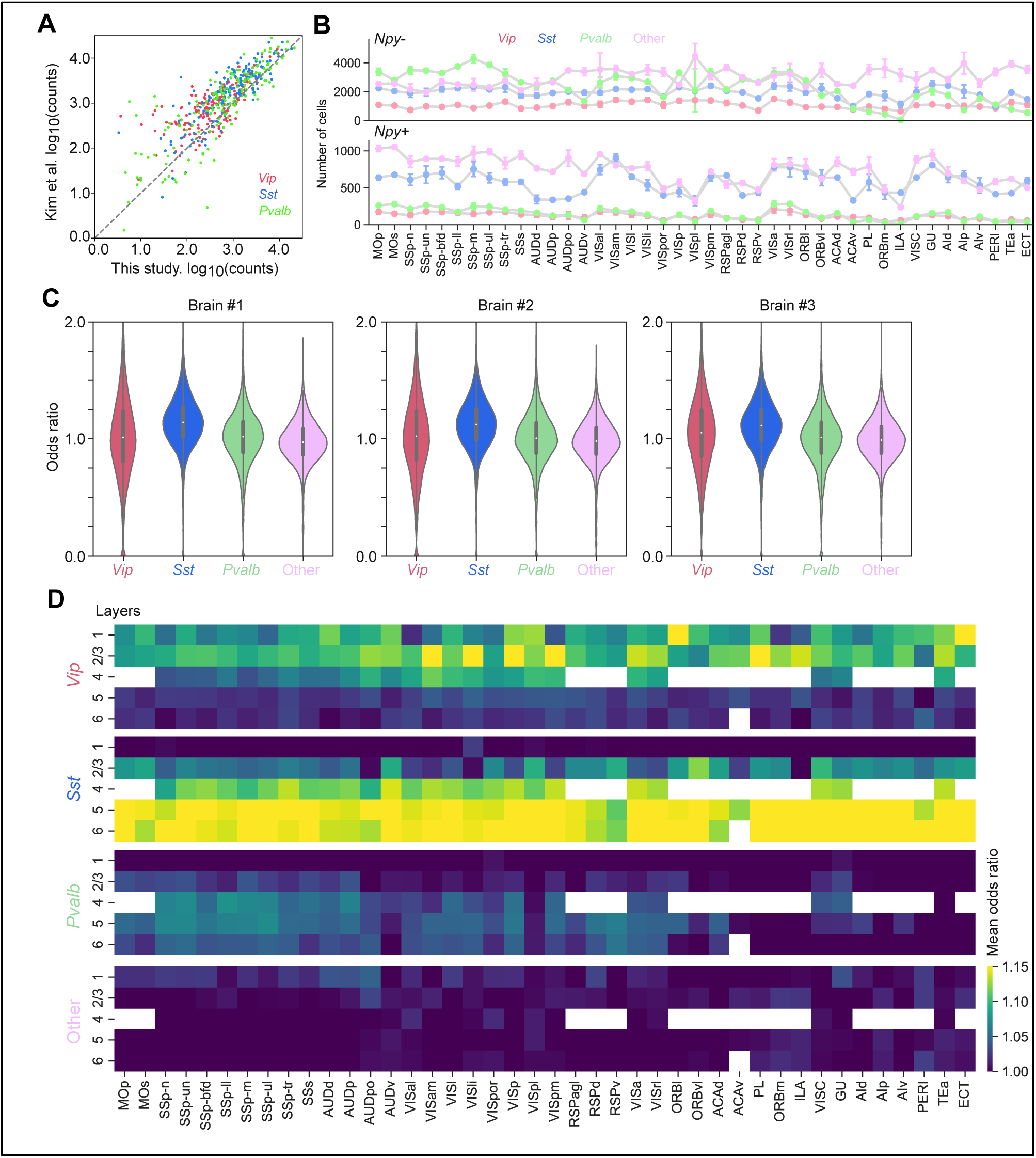
Cortex-wide mapping and interaction analysis of inhibitory neurons and oligodendrocytes. (**A**) Correlation plot of the regional number of the inhibitory neurons between this study and the report of Kim et al., 2017. Each dot indicates the anatomical region. Both of the annotations of anatomical regions came from Allen Brain Atlas. (**B**) The *Gad1*+ cells were classified into four groups (*Vip*+, *Sst*+, *Pvalb*+, and Other). The four groups were further classified based on the expression of *Npy* and the cell numbers in each anatomical region are shown (N=3). (**C**) Violin plot of the odds ratio of the candidate satellite oligodendrocytes. The replicates in three brains are shown. (**D**) The heatmap of the odds ratio based on the anatomical regions and cortical layers. The mean values of the odds ratios are visualized.

**Figure S4.**
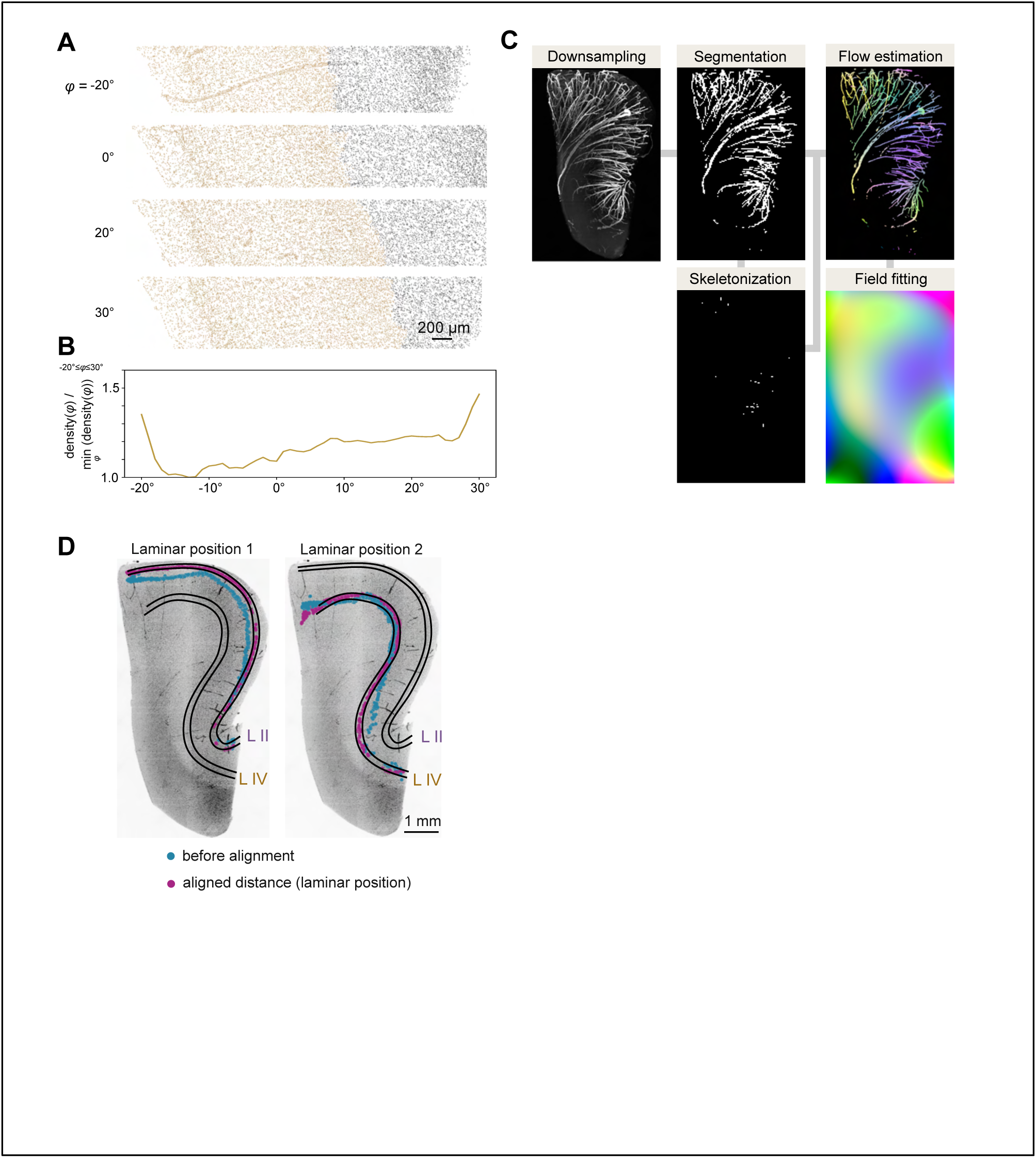
Reconstruction of the morphogenic flow from arteries. (**A**) The reconstructed virtual tissue slices from 3D volumes for the top of Figure 4B. Each dot indicates the position of a nucleus. The cells in layers I∼IV used for the analysis are shown in color. (**B**) The densities (cell number per volume) were calculated and plotted. The sharp transition at -20° was probably due to the thick artery. We can see the variability in cell densities by pivoting the sectioning angles. The values normalized by the minimum density are shown. (**C**) The workflow to reconstruct the vector field of morphogenic frow from the signal of arteries. See also **Supplementary Methods**. (**D**) The tracks on a specific laminar position are plotted. The tracks before the alignment are also plotted for comparison. The left plot indicates that the laminar position overlaps with the manual annotation of layer II. The right plot indicates that the laminar position overlaps with the manual annotation of layer IV. The maximum intensity projection of 200-μm-thick nuclear-stained tissue is overlaid. L: layer.

**Figure S5.**
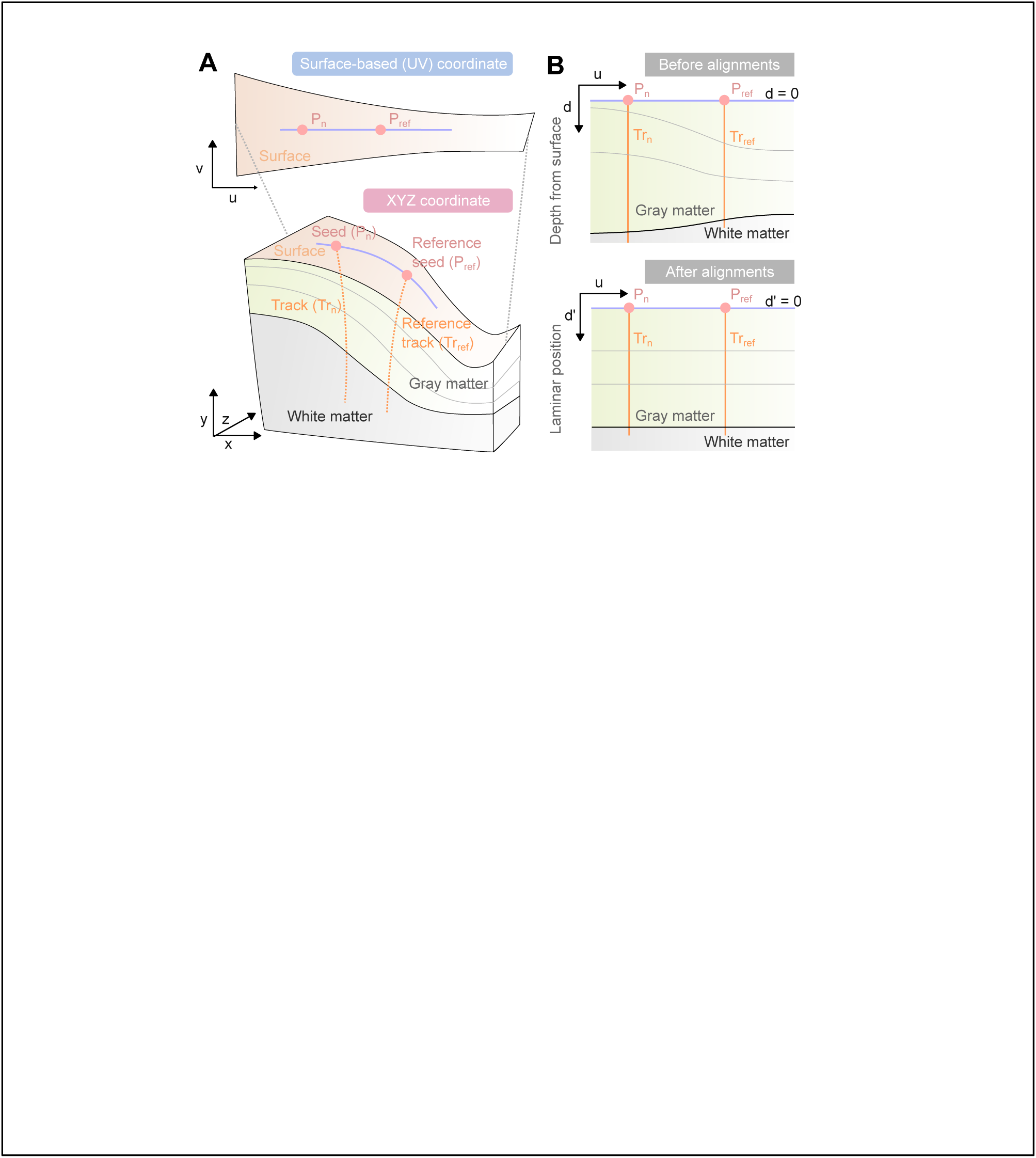
Illustration of the surface-based coordinates and alignments of morphogenic tracks. (**A**) The surface of the cortex in the original space in the microscope (XYZ coordinate) was non-linearly transformed into a two-dimensional coordinate (surface-based coordinate). See also **Supplementary Methods**. The cortical flattening was used for visualization purposes. (**B**) The alignment of the morphogenic tracks. The depth from the surface (*d*) was transformed into the laminar position (*d*’) so that comparisons of tracks are possible.

**Figure S6.**
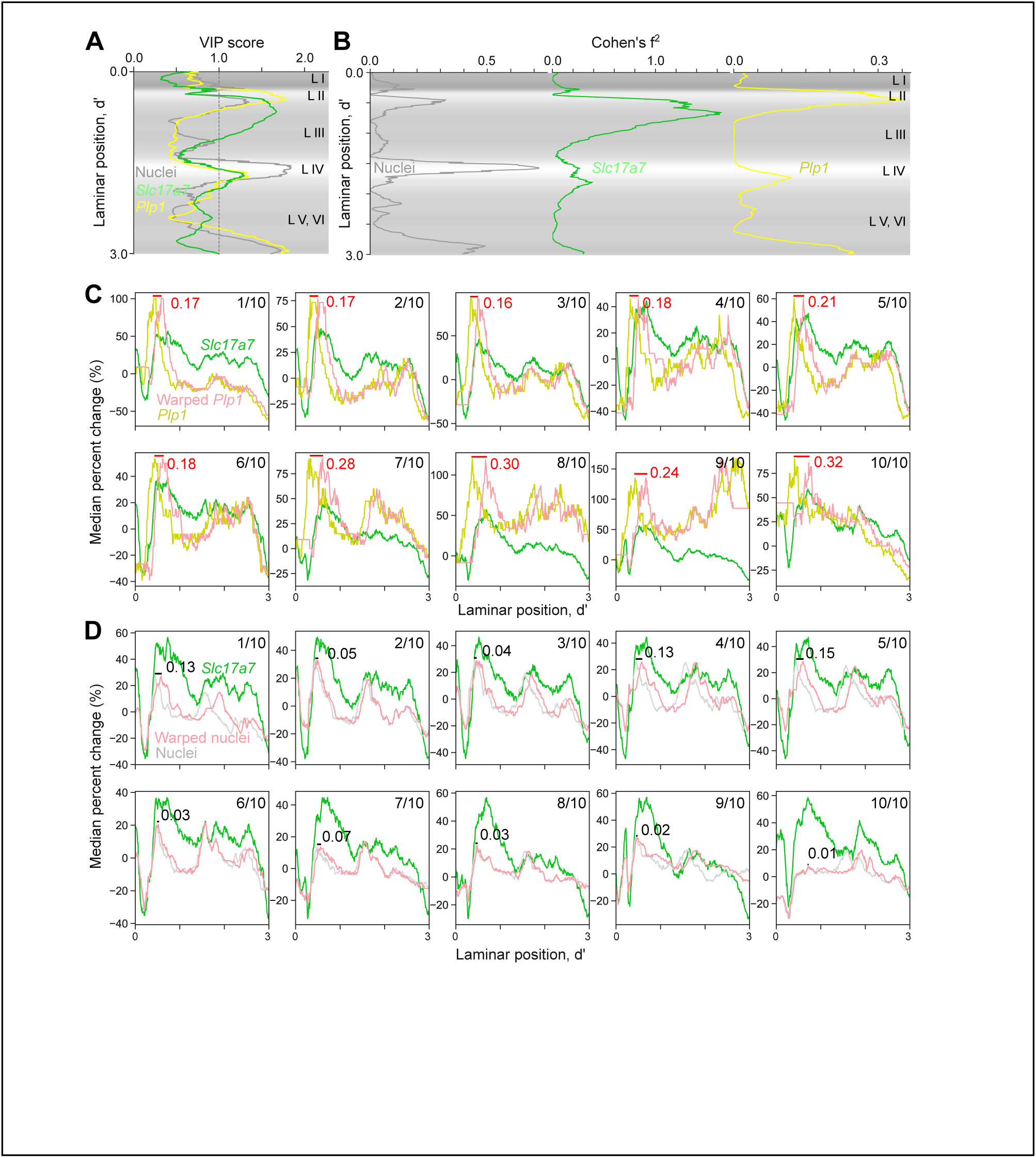
Synchronic emergence of excitatory neurons and oligodendrocytes. (**A**) VIP score of PLS-DA by cell types. The averaged cell density with a gray color code is shown in the background. (**B**) Cohen’s f^2^ at each laminar position. We applied a multiple linear regression model with a label of the region and local flux as covariates to predict the cell density at the laminar position. The averaged cell density with a gray color code is shown in the background. (**C**) The quantification of the peak shifts between profiles of median percent changes of *Slc17a7*+ cells and *Plp1*+ cells. The tracks were classified into 10 groups based on the percentile of the local flux. The group is indicated at the top right corner. For example, 2/10 indicates the group of the tracks with the local flux in the range of 10 to 20 percentiles. The profiles were aligned in the same way we did in the alignments of the morphogenic tracks. The warped profile is shown in red. The peak of the original profile and the warped profile was compared and the distance is shown in each plot. (**D**) Quantification of peak shifts between profiles of median percent changes of *Slc17a7*+ cells and nuclei. The peak shift analysis was performed in the same way as **C**. L: layer.

